# A growth chart of brain function from infancy to adolescence based on electroencephalography

**DOI:** 10.1101/2023.07.07.548062

**Authors:** Kartik K. Iyer, James A. Roberts, Michaela Waak, Simon J. Vogrin, Ajay Kevat, Jasneek Chawla, Leena M. Haataja, Leena Lauronen, Sampsa Vanhatalo, Nathan J Stevenson

## Abstract

**Background:** In children, objective, quantitative tools that determine functional neurodevelopment are scarce and rarely scalable for clinical use. Direct recordings of cortical activity using routinely acquired electroencephalography (EEG) offer reliable measures of brain function.

**Methods:** We developed and validated a measure of functional brain age (FBA) using a residual neural network-based interpretation of the paediatric EEG. In this cross-sectional study, we included 1056 children with typical development ranging in age from 1 month to 18 years. We analyzed a 10 to 15 minute segment of 18-channel EEG recorded during light sleep (N1 and N2 states).

**Findings:** The FBA obtained from EEG had a weighted mean absolute error (wMAE) of 0.85 years (95%CI: 0.69-1.02; n = 1056). A two-channel version of the FBA had a wMAE of 1.51 years (95%CI: 1.30-1.73; n = 1056) and was validated on an independent set of EEG recordings (wMAE = 2.27 years, 95%CI: 1.90-2.65; n = 723). Group-level maturational delays were also detected in a small cohort of children with Trisomy 21 (Cohen’s *d* = 0.36, *p* = 0.028).

**Interpretation:** An FBA, based on EEG, is an accurate, practical and scalable automated tool to track brain function maturation throughout childhood with accuracy comparable to widely used physical growth charts.

**Funding:** This research was supported by the National Health and Medical Research Council, Australia, Helsinki University Diagnostic Center Research Funds, Finnish Academy, Finnish Paediatric Foundation, and Sigrid Juselius Foundation.

**RESEARCH IN CONTEXT:** *Evidence before this study:* Tools for objectively tracking neurodevelopment in paediatric populations using direct measurement of the brain are rare. Prior to conducting this study, we explored multiple databases (Google Scholar, PubMed, Web of Science) with search strategies that combined one or more of the terms “paediatric brain development”, “brain age”, “age estimation”, “MRI measurements”, “EEG measurements”, “machine learning”, “artificial intelligence”, “advanced ageing”, “neurodevelopmental delays” and “growth charts” with no restrictions on language and dates. In screening over 500 publications, 7 studies evaluated brain age in children using MRI and only a single study investigated maturation in EEG activity across discrete age bins.

*Added value of this study:* We formulated a measure of functional brain age (FBA) using state-of-the-art machine learning (ML) algorithms trained on a large, unique database consisting of multichannel clinical EEG recorded from N1/N2 sleep (n = 1056 children; 1 month to 17 years), with typical neurodevelopment confirmed at a 4-year follow-up. The FBA showed a high correlation with age and detected group-level differences associated with conditions of neurodevelopmental delay.

*Implications of all the available evidence:* Age is prominent within EEG recordings of N1/N2 sleep and is readily extracted using ML. Public release of the FBA estimator and the use of EEG, commonly delivered in outpatient settings, as the basis of age prediction enables clear translation of measures of ‘brain age’ to the clinic. Future work on EEG datasets across various neurodevelopmental profiles will enhance generalisability and user confidence in the clinical application of brain age.

## INTRODUCTION

Deriving brain age in relation to a person’s chronological age has become a burgeoning focus for scientists, clinicians, and education providers.^1^ The physiological maturation of the brain is fundamentally shaped by an interaction between an individual’s genetic traits and acquired/environmental effects.^2,3^ The chronological age of an individual is a particularly important benchmark in the assessment of paediatric populations, given that over 10% of children worldwide^4^ are affected by neurodevelopmental problems.

Current predictors of brain age are built predominantly upon metrics of structural MRI morphometry such as cortical thickness, grey matter, white matter, and intracranial volumes^2,3,5,6,7^ derived from large bio-banks of magnetic resonance imaging (MRI) scans. Across these studies, the key biomarker is defined as the difference between predicted brain age and chronological age.^8^ Brain predicted age difference (PAD) has been associated with neurodegeneration and compromised neurological health in adults^9,10^ and autism spectrum disorder in children.^11^

We propose the use of electroencephalography (EEG) as a basis of age prediction. The EEG’s high temporal resolution enables the capture of subtle changes in neurophysiological function across dynamic brain states such as evoked responses, resting states and sleep providing valuable insights into typical and atypical maturation during neurodevelopment.^12,13,14,15^ Chromosomic and genetic alterations, such as those present in children diagnosed with Trisomy 21, Autism spectrum disorder or attention-deficit/hyperactivity disorder, affect brain function at the neuronal level.^16,17,18^ Developing neurodevelopmental biomarkers that are associated with these cellular-level changes to brain functions could thus provide key actionable indicators that guide early intervention and personalisation of clinical care to ultimately impact long term outcomes.

Brain age predictors that track maturation, as measured by the EEG, are promising in paediatrics due to rapid functional changes that occur in concert with, or in parallel to, structural changes in the brain during childhood and adolescence^19,20,21,22^ unlike structural MRI which shows promise in brain age prediction but only captures spatial information. EEG (and fMRI) captures neurophysiological activity across space and time. Brain age prediction via these functional modalities is emerging^23,24,25,26,27^, but is currently limited to short acquisitions in controlled research settings or incomplete representation of the entire paediatric age range.^2,3,6,7,8,28^ Establishing a “functional brain age” (FBA) in paediatric cohorts can, therefore, complement the array of behavioral assessments typically employed in clinical practice, enhancing the assessment of neurodevelopment.

In this study, we charted the growth of brain function using an EEG-derived FBA. Machine learning methods applied to a large cohort of EEG recordings from children with typical development formed the basis of the FBA. We used light sleep (N1 and N2) due to the ubiquity and comparability of these neurophysiological states across childhood and tested the diagnostic potential of the FBA using a small cohort of children with atypical neurodevelopment.

## METHODS

### Study Design

The framework for predicting functional brain age (FBA) from sleep EEG is presented in **Figure 1**. A FBA was developed using a residual neural network architecture (Res-NN). A ‘bag of features’ and Gaussian process regression (GPR) predictor of age was used to benchmark age methods. Our input for the FBA model consisted of 60 second epochs of EEG, where an FBA per recording was calculated as an average across multiple epochs extracted from a 10 or 15 minute segment. We developed the FBA on a primary training dataset (D1) which comprised 15 minutes of 18-channel EEGs from 1056 children recorded at the Helsinki University Children’s Hospital, Finland. We then validated our trained model on a dataset (D2) which comprised of 10 minutes of 2-channel EEGs recorded from 723 children at Queensland Children’s Hospital, Brisbane, Australia.

**Figure 1.**
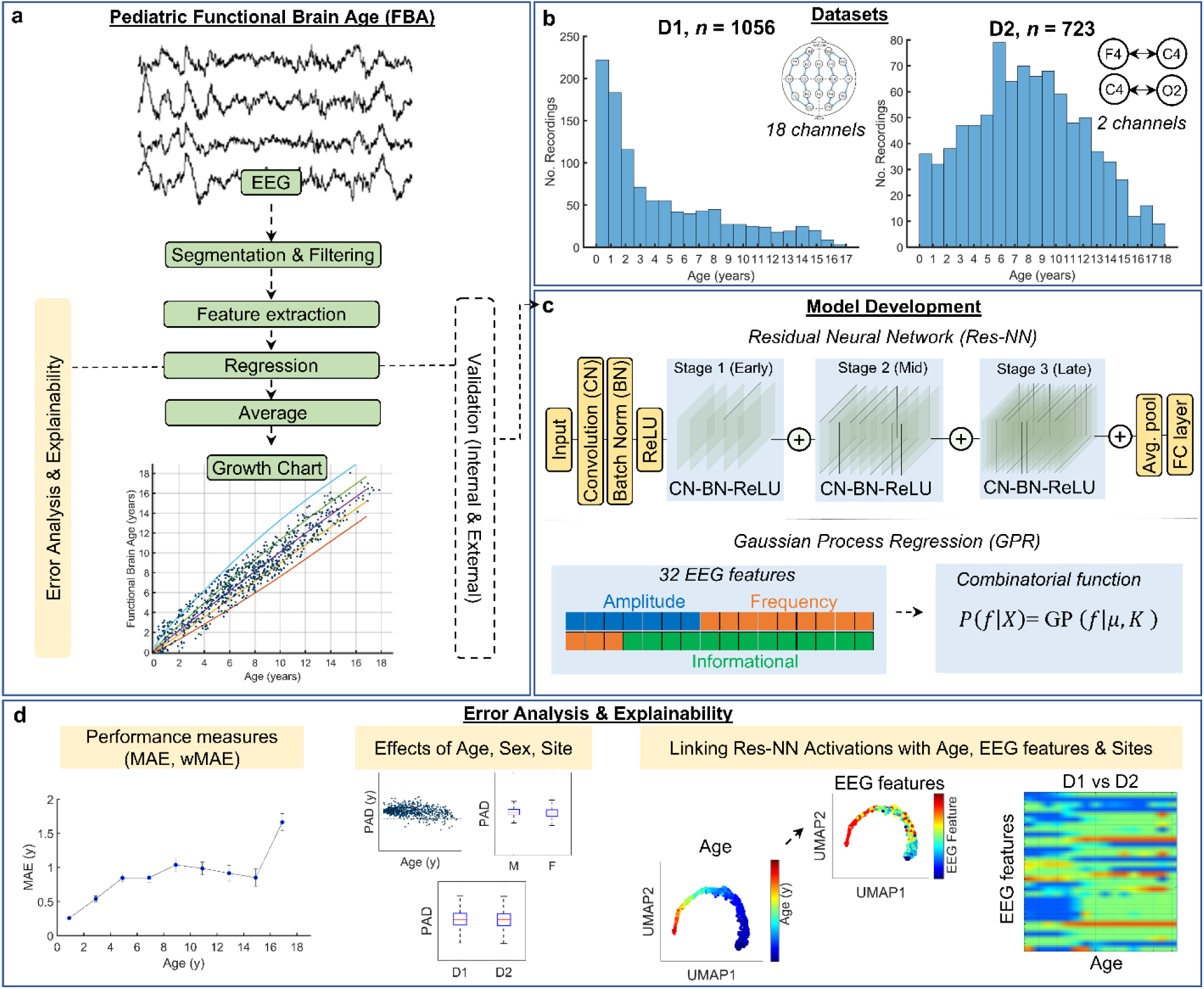
Study design. **a.** EEG acquisition from routine sleep studies from 2 sites (D1 – Helsinki, n = 1056; and D2 – Brisbane; n = 723). The development of a functional brain age (FBA) growth chart within a supervised learning framework. Single, 60 s EEG epochs per recording were used to form a preliminary estimate of FBA that was then averaged (median operation) across all available epochs within a 10-15 minute segment of EEG. Model evaluation (FBA) involved cross-validation procedures from the primary developmental dataset (D1) with external validation on an independently collected dataset (D2). Growth charts for D1, D2 and a combined D1+D2 dataset were computed. **b.** Distributions for training (D1) and validation (D2) datasets across age and the EEG channel montages of acquisition. Data consisted of 5 minute epochs of N1 sleep followed by 10 minute epochs of N2 sleep from 18 channels for D1, and 10 minute epochs of N2 sleep from 2 channels for D2. **c.** A trained Residual Neural Network (Res-NN) was our primary method of feature extraction for FBA prediction. Performance was benchmarked against a GPR model on *a priori* engineered EEG summary measures.^29^ **d**. Performance of the FBA was assessed via measures including the mean and weighted absolute error (MAE, wMAE) and predicted age differences (PAD = FBA minus chronological age). Effects of age, sex, and recording site were examined using statistical tests. The behaviour of the final trained Res-NN was explained using links between network activation, age, and engineered EEG features across sites.

### Datasets

Our primary training dataset (referred hereto as *D1*) was collected from a convenience sample of EEGs collected at the Helsinki University Children’s Hospital in Helsinki, Finland. EEG was recorded using 10-20 electrode positions with an electrode on the vertex as an active reference (either Fz or Cz) using a Nicolet One EEG (Natus Medical Inc. Middleton, WI, USA). All EEG recordings were sampled at 250 Hz and the referential montage was saved in a pseudonymised EDF file format. A total of 19 channels (Fp1, Fp2, F7, F3, Fz, F4, F8, T3, C3, Cz, C4, T4, T5, P3, Pz, P4, T6, O1, O2) were collected as part of the EEG; reference EEG electrodes were attached to the mastoid (A1/M1 and A2/M2). All children (aged between 1 month to 18 years) with an EEG recorded between 2011 and 2016 were screened. EEG recordings in D1 were clinically reviewed to define normality of the record, where the montage of review was a standard longitudinal bipolar montage (double banana). The EEG record for D1 was re-reviewed by EEG technicians trained for the purpose (and approved by L.L).

A total of 1056 children with typical neurodevelopment were available for analysis across the 18 year age range (see **Supplementary Table 1** for demographics and dataset comparisons). For D1, the reporting of sex was derived from the Finnish social security system where sex is medically defined. A 15 minute segment of EEG was extracted and saved in EDF format. The first 5 minutes of each segment consisted of N1 sleep and the remaining 10 minutes consisted of N2 sleep. The transition between N1 and N2 sleep was defined by the first occurrence of sleep spindles or K- complexes^30^ (sleep was scored using a referential montage). These 15 minute segments may include very brief (paroxysmal) arousals^31^ that may typically disrupt physiological sleep in children but were not seen to corrupt the EEG due to their transient nature. Before undergoing a clinical EEG session, families were asked to wake up their child 2 to 4 hours earlier than their usual wake-up time, to ensure they would be able to fall asleep in the laboratory; however, individual variations in sleep pressure were not measured. This early period of EEG comprising N1/N2 transitions was selected for FBA training.

Our external validation dataset (referred hereto as *D2*) was collected from a convenience sample of polysomnography (PSG) studies from the Queensland Children’s Hospital (QCH) in Brisbane, Australia (Respiratory and Sleep Medicine Department), reviewed between 2014 and 2021. PSG was acquired via the EMBLA N7000 (Natus Neuro, Middleton, WI, USA). For D2, a total of 3 channels were recorded overnight (F4, C4 and O2) as part of the PSG. EEG was recorded using 10-20 electrode positions and recordings were sampled at 200 Hz or 500 Hz; reference EEG electrodes were attached to the mastoid (A1/M1 and A2/M2).

Following screening of D2 data, a total of 723 children with typical neurodevelopment were available for analysis across the 18 year age range based on normal outcomes following PSG review (see **Supplementary Table 1** for demographics and dataset comparisons). We also identified a cohort of children with Trisomy 21 (*n* = 40) in D2 whom had normal outcomes following PSG review, to examine group-wise differences in FBA. For all D2 data, the reporting of sex was obtained from the Queensland Health record, which is determined by the parent or guardian of the child at the initial referral and visit to the Public Health Service.

As per D1, all sleep stages were seen and scored by a clinician according to the American Academy of Sleep Medicine (AASM) guidelines using a referential montage. For D2, we limited our EEG analysis to the first 10 minute period of N2 sleep only due to limited availability of N1 in D2 data (N1 was only present in 45/723 PSGs). The age of children across both D1 and D2 was resolved in months with **Figure 2** summarising the screening flowchart for these datasets.

**Figure 2.**
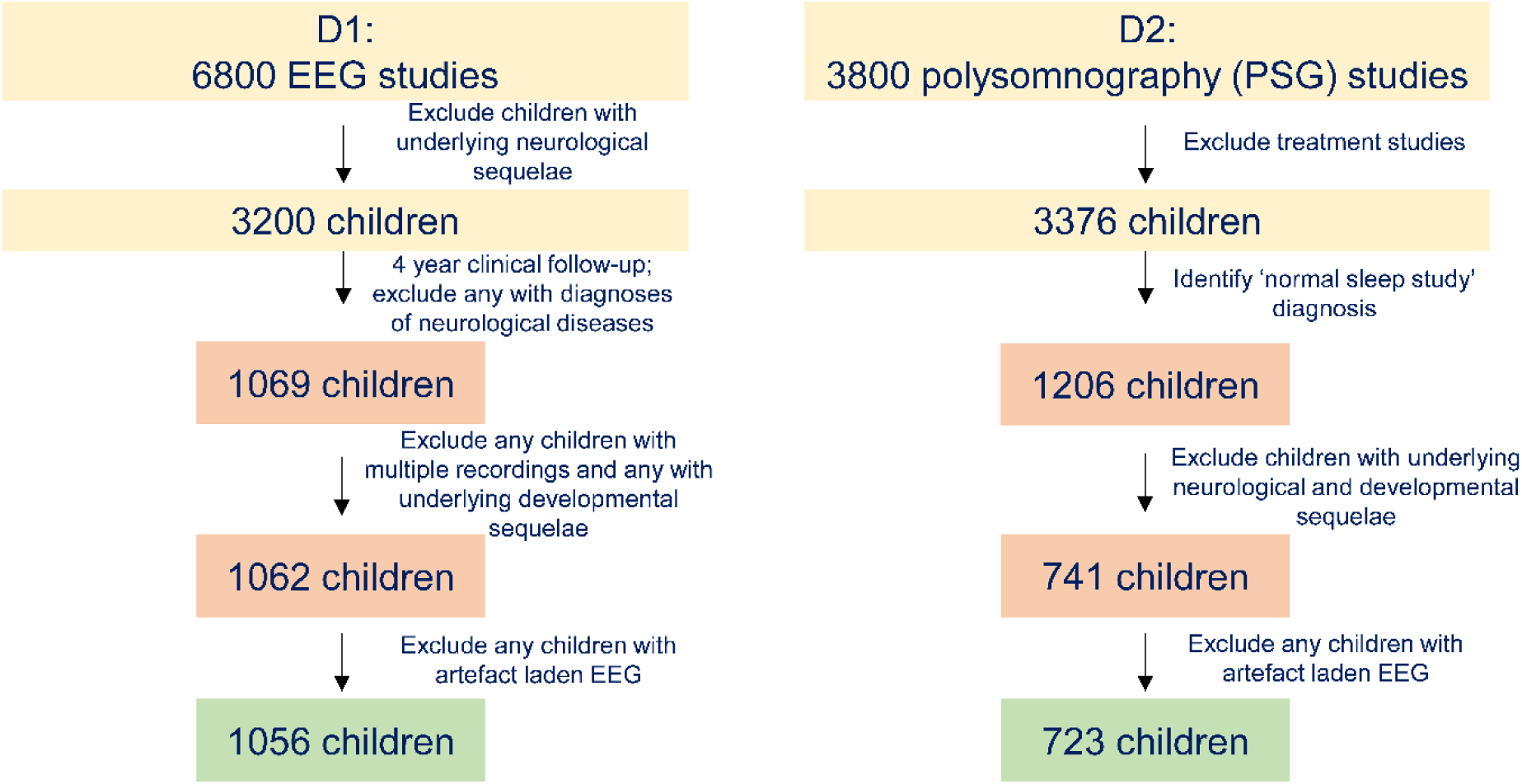
Screening flowchart for datasets used to train functional brain age algorithms. For D1, 1056 children with an additional 4 year clinical follow-up were included, which enabled us to identify any further neurological diagnoses that may exclude them from further analyses. A final technical check was performed to identify outliers due to significant artefacts, some of which could be removed by re-export of the EDF format. For D2, 723 children with a diagnostic label of ‘normal sleep study’ and no underlying neurodevelopmental diagnoses were included.

### Processing the EEG with machine learning tools

#### Data processing

All EEG data were zero-phase filtered in both forward and reverse directions with an infinite impulse response, bandpass, 4^th^ order Butterworth filter with a lower cutoff of 0.5 Hz and an upper cutoff of 30 Hz (GPR) or 15 Hz (Res-NN). EEG data were resampled to 64 Hz and 32 Hz as inputs to the feature extraction components of the GPR and residual network regression, respectively. Residual network regression approaches employed EEG data inputs at a lower sampling rate to reduce the size of training data and maximise computational efficiency, with several qualitative tests performed to ensure that important aspects of sleep EEG were retained (e.g. delta and alpha rhythms in sleep). Our feature-based methods were examined at a higher sampling frequency and ensured that higher frequency components of the EEG were also captured across age. For D1, a bipolar montage was computed from the monopolar/referential EEG data resulting in 18 EEG derivations/channels: Fp2-F4, F4-C4, C4-P4, P4-O2, Fp1-F3, F3-C3, C3- P3, P3-O1, Fp2-F8, F8-T4, T4-T6, T6-O2, Fp1-F7, F7-T3, T3-T5, T5-O1, Fz-Cz, and Cz-Pz.

Further, each 15 minute segment of EEG was divided into 60 s epochs for analysis. Epochs were extracted with a 30 s overlap (29 epochs per recording). We assumed that 60 s was sufficiently long to capture important EEG signal characteristics and short to reduce any effects of non-stationarity in the EEG while generating a sufficiently large and diverse set for model training.

For D2, a simplified bipolar derivation of the EEG; i.e., F4–C4, C4–O2 was used due to the limited availability of channels. Each 10 minute EEG recording in D2 was segmented into 60 s epochs with a 30 s overlap (19 epochs per recording) and used for training and testing.

At the end of these data curation steps, we then developed a residual neural network regression (Res-NN) for age prediction.^32^ We also used GPR model as a benchmark.^33^

#### Res-NN

EEG epochs were first resampled to 32 Hz (resampling included anti-aliasing filtering). We added variability to the residual neural network by changing the temporal filter width (FW), filter channel depth (FD) and filter number (FN) within the convolutional layers as well as increasing the network depth (ND). We used the file *generate_networks_v2.m* to generate networks with different configurations and architectures (see **Supplementary Figure 1**; code provided in our GitHub repository, details in *Data sharing statement*).

Several parameters specific to the definition of these neural network architectures were selected during each training iteration. In general, parameters defined the filter size (temporal width and channel depth), filter number and network depth. Training options such as solver type and mini-batch size were selected based on preliminary analysis (see **Supplementary Figures 2 to 9**), and an alternate architecture based on inception layers was also evaluated.^34^

#### GPR

This approach combined several summary measures of the EEG features to form a prediction of age. We chose GPR due to its consistent performance across various imaging modalities in estimating brain age following feature extraction due to its capability to model the underlying latent distribution and quantifying any associated uncertainty to provide probabilistic predictions from data. ^2,3,35,36,37^ All EEG epochs were resampled to 64 Hz (resampling included anti-aliasing filtering). We used a total of 32 features from the EEG. Of these, 7 features were general measures of EEG amplitude activity, 12 features were frequency dependent representations present within the EEG, and the remaining 13 features were further computations of information content drawn from amplitude, frequency, and entropy based transformations of the EEG signal. A descriptor of the EEG features used is listed in **Supplementary Table 2** and shown **in Supplementary Figure 10**; EEG feature extractor available in our GitHub repository, details in *Data sharing statement*). For both D1 and D2, the set of 32 EEG features were estimated for each available channel and then averaged across channels (with a median operation used). For training, each EEG recording was, therefore, summarised by a C-by-M-by-32 feature matrix (where C was the number of channels, M was 29 epochs per recording for D1 and 19 epochs per recording for D2, and there were 32 features per epoch). For D1, the full feature set for 1056 children thus resulted in 18 channels by 29 epochs by 32 features, which were then averaged (median) across channels and epochs resulting in a 1056 by 32 feature matrix. The same procedure was applied to the reduced channel montage of D1 (i.e. 2 channel montage) and for D2 EEG data, which resulted in a 723 by 32 feature matrix. Similar to the Res-NN, we expected that temporal averaging of FBA estimates per child across all available epochs. A combinatorial function (shown in **Figure 1**) was used to determine the final brain age estimate. This was achieved by first assuming a Gaussian prior (GP) where *P*(*f* |*X*) = GP(*f*|*μ*, *K*) denotes the dependence of *f* (i.e. FBA) on *X* (i.e. age) across all observed data points and µ and *K* represent the mean function and kernel parameters, respectively.

#### Performance assessment, cross-validation, optimisation, and independent validation

##### Performance assessment

We utilised two commonly used measures to compare the predicted age to actual age and, therefore, define the accuracy of prediction: mean absolute error and root mean square error. The root mean square error (RMSE, Eq.1) between predicted age per recording and actual age was defined as

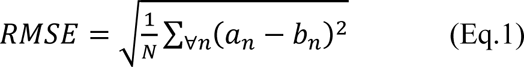

where *a*_*n*_ is the age of the *n*^th^ EEG recording in years, *b*_*n*_ is the median predicted age across all extracted EEG epochs per recording, and *N* is the number of EEG recordings (1056 for D1 and 723 for D2). The mean absolute error (MAE, Eq.2) was defined as:

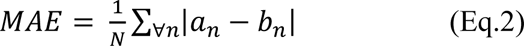

We supported these performance measures with weighted MAE. The wMAE is a variation of the MAE that normalises across the distribution of the data with respect to age; i.e., an approximation of the MAE for a cohort with uniformly distributed age. Based on the age distribution and sample sizes of our cohorts, we defined it as the average MAE across age binned averages of the MAE, specifying a bin width = 2 years, ensuring that enough samples were represented in each age bin. We also computed the relative error with respect to age, by calculating the percentage change between predicted age (FBA) and chronological age.

##### Cross-validation, optimisation and training

We used D1 as our primary training set for developing an FBA. To evaluate the accuracy of prediction, we used 10-fold cross-validation where the train/test split of EEG recordings was 90% for training and 10% for testing. This process was repeated 10 times so that all EEG recordings were excluded from training for at least one iteration (fold).

For Res-NN, due to computational limits, initial experiments of randomly selected combinations of network architectures were used to select several model parameters based on maximum regression accuracy (minimum RMSE) on 10-fold cross-validation outputs. These parameters included training solver and mini-batch size. It was assumed that the selection of these parameters was less prone to overfitting than internal network weights from variability in architecture. Architecture parameters were selected using an internal 10-fold cross-validation (9:1 training data split). We used a small fixed grid approach and selected the parameter combination (filter width, filter depth, filter number, network depth) that minimised the RMSE. A single optimal architecture was defined by averaging internal validation RMSE across all 10 training iterations (a point of data leakage assumed to result in negligible overfitting). Remaining training hyperparameters were not optimised: initial learning rate was 0.0001 for training for initial selection and nested architecture selection, training drop factor was 0.1 every 8 epochs, maximum number of training iterations was 50, 10% of recordings within the training set were used for internal validation, and if no changes in RMSE on the validation set were detected within 6 training iterations stopped training. The squared gradient decay factor was 0.99 for the ADAM solver. Data extraction and network implementation and training was performed in MATLAB (The MathWorks Inc, Natick, Massachusetts, USA: Deep Learning Toolbox; R2020a or R2021a with GTX 1070 or 1080 graphics cards, for dataset D1 and datasets D2 and D1+D2, respectively). On average, training time per fold for D1 with an 18 channel bipolar montage was 16.4 minutes; for 2-channel montages for either D1 or D2 training time per fold on average was 4 minutes.

To explore how the EEG signal was represented by the neural network, we extracted activations of network layers at several stages of the deep learning network when trained and tested on all available data (for this exploratory analysis no cross-validation was used). Here, the optimal network was composed of 63 layers, where an early layer output was taken at layer 6, the middle layer output was taken at layer 29 and the late layer output was extracted at layer 61. A Uniform Manifold Approximation and Project method (UMAP^38^) was used to reduce the high-dimensional network activation space into a lower dimensional 2D space (UMAP1, UMAP2), wherein the output space was coded according to age. Visualisations via UMAP allowed us to qualitatively assess how the activations of a network layer cluster with respect to age (see **Figure 4b**). To quantify the association between UMAP1 and UMAP2 dimensions and age, we used GPR with UMAP1 and UMAP2 as inputs to predict age (via 10-fold cross-validation). The accuracy of age prediction for each network layer was then derived. We used the same approach to track how UMAP representations of network activations were linked to individual features of the EEG (**Supplementary Figure 11**).

**Figure 3.**
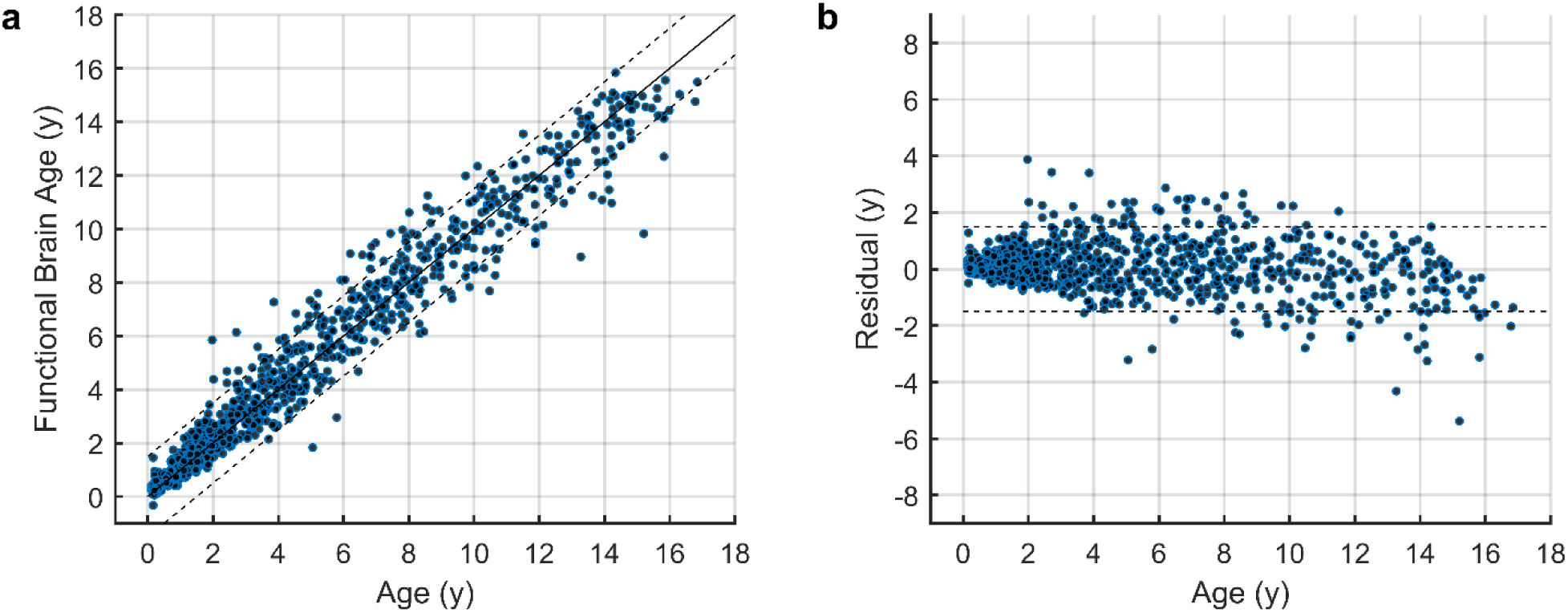
Functional brain age estimated in children and performance in training dataset (D1, n = 1056). **a.** Res-NN model (R^2^=0.96, MAE=0.56 years). The dashed line indicates an error bound of ±1.5 years. Individual EEG recordings (blue filled circles) are plotted across ages. **b.** The residual error (predicted age difference; PAD) represents the difference between the FBA measure and chronological age of an individual, in years.

**Figure 4.**
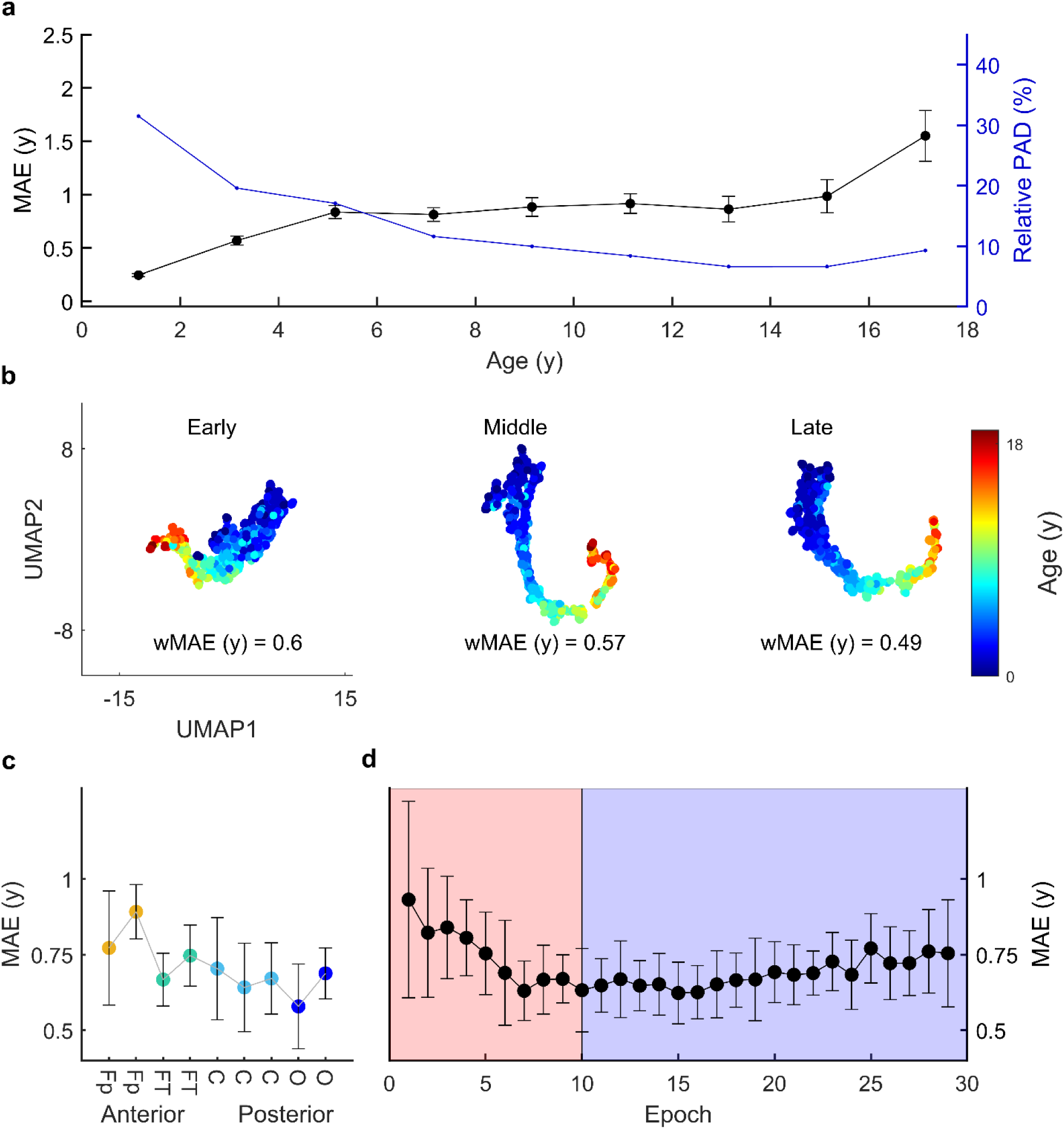
Interpreting the EEG-based FBA. **a.** The mean absolute error (MAE) of the Res-NN (black circles, left axis) across 2 yearly age bins and the relative accuracy of PAD (in blue, right axis) across 2 yearly age bins. For both plots, the mean and standard error of the mean (SEM) are plotted to reflect the sample distribution within age bins. **b.** A lower-dimensional representation of the Res-NN network generated by UMAP on 18-channel D1 data. The network is composed of 63 layers represented by early (layer 6), middle (layer 29) and late stages (layer 61). Here, the Res-NN of the EEG clusters into younger age groups (blue) and older age groups (red) throughout the training phase, wMAE was calculated using UMAP values as predictors (no cross-validation). **c.** The performance accuracy of the Res-NN model (MAE, years) following individual cross-validation per EEG channels. Colors are ordered by anterior to posterior channel derivations, with frontopolar (Fp, in yellow), frontotemporal (FT, in teal), central (C in light blue) and occipital channels (in dark blue). Average MAEs are shown with the standard deviation shown as error bars. **d.** The temporal change in MAE of epoch sequences obtained sequentially from N1 (light pink) and N2 (light purple) indicated that the lowest MAEs were observed during a transition between sleep N1 and N2 stages. The MAE is shown with error bars indicating the standard deviation.

For GPR, hyperparameters were selected using Bayesian optimisation within a nested cross-validation and included kernel function (e.g. rational quadratic) and sigma values. Shapley values ^39^ were also computed to quantify each feature in terms of its contribution to the overall prediction of FBA (**Figure 5c**), extending on the notion of linear model predictions.

**Figure 5.**
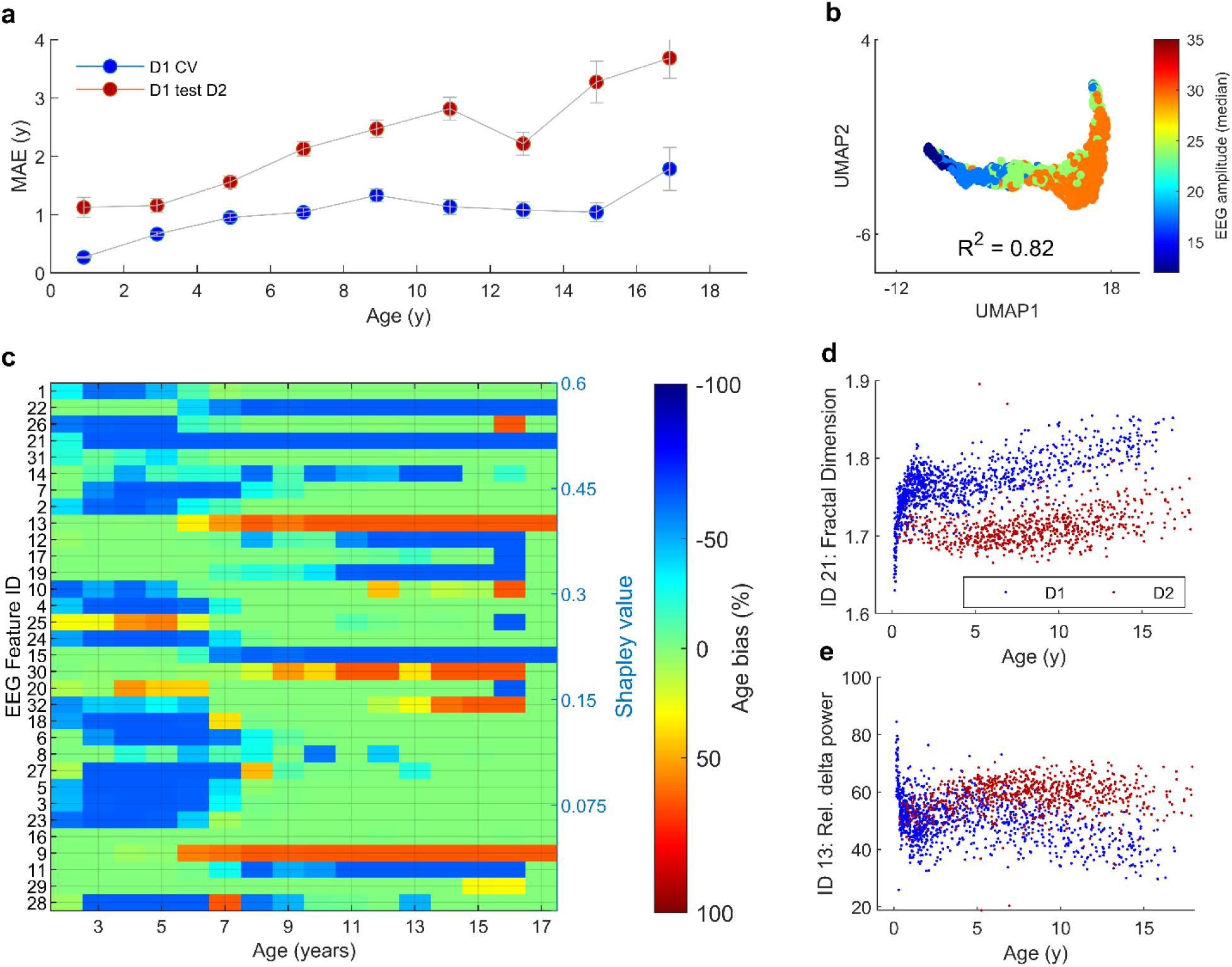
Validation of the FBA across sites. **a.** Differences in MAE between D1 (n = 1056) and D2: cross-validation (train and test D1 CrossVal, indicated by blue circles) versus external validation on D2 (train D1 test D2, red circles). The MAE across age is higher when testing D1’s model on D2 (n = 723). The MAE is shown with error bars indicating the standard deviation. **b.** Mapping EEG features onto the late layer network UMAP (layer 61) derived from 2-channel D1 data; here we show the EEG median amplitude as an exemplar. The R2 value indicates the strength of correlation between UMAP values and feature value, based on a GPR prediction. **c.** Differences between EEG features and site across age. EEG features (IDs 1 to 32, see also Supplementary Table 2 for feature names) were ordered by Shapley values (highest to lowest) to indicate the relative contribution of the feature to the overall model. Red and blue colors indicate significant age biases following multiple comparisons correction (Bonferroni’s method). A positive age bias percentage (red) indicates EEG feature values being ‘older’ in D2 compared to D1 and a negative age bias percentage (blue) indicates EEG feature values being ‘younger’ in D2 compared to D1. **d.** Fractal dimension (feature ID 21) and relative delta power (0.5 to 2 Hz, feature ID 13) are exemplar EEG features that show site related differences (D1=blue; D2=maroon).

### Statistics

#### Inclusion/exclusion criteria

We included EEG recordings of children who exhibited normal ranges of background activity for their age and met the criteria of being typically developing based on neurodevelopmental and/or physical outcomes, including confirmation of typical development at a four-year clinical follow up. Children with EEGs showing seizure-related or aberrant paroxysmal discharges were excluded. Incomplete data caused by technical difficulties were also excluded. Additionally, children with diagnoses of neurological diseases (including epilepsy), sleep disorders, psychiatric disorders, brain-acting medications, presence of tumors/cancer, congenital and/or perinatal disorders, and malformations were excluded. Children with neurodevelopmental and/or neurodegenerative conditions such as Autism spectrum disorder, Duchenne Muscular dystrophy, Spina bifida, Spinal muscular atrophy, Trisomy 21 were excluded. Lastly, children with seizure activity or central sleep apnea events higher than 5 events per hour were also excluded. If multiple recordings were collected in the same children, only one time point was selected based on age distribution of the larger cohort. Therefore, each child in D1 (and D2) only had one EEG recording that was subsequently used for analysis. Lastly, EEGs with significant artefact identified via visual review and computational analysis of the data (e.g. excessively high mean amplitudes, kurtosis of amplitude envelope, and Hjorth parameters for age) were also excluded.

As both datasets originated from large convenience samples, sample size calculations were not performed.

#### FBA accuracy testing

We first evaluated the prediction accuracy of FBA between Res-NN and GPR models. The key variable used in this analysis was the residual error — also referred to as the predicted age difference, PAD (the difference in years between the FBA and chronological age). When comparing between age estimators (Res-NN, GPR) with various feature subsets and mean cohort age), we used a t-test for paired samples with absolute PAD as the input where our null hypothesis was that the absolute PAD would not be different between Res-NN and comparative estimator (as we used simulated data to evaluate an estimator based on head circumference we use a t-test for unpaired samples). We also estimated effect size using Cohen’s D (for paired and unpaired samples as necessary).

We then analyzed potential confounding effects of age and sex on PAD using multiple linear regression. Here, our multiple regression models for D1 and D2 tested potential effects of sex and age (Eq.3).

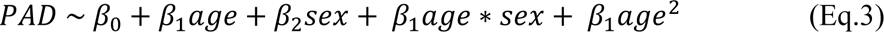

We assessed the performance of the FBA across spatial and temporal combinations of the EEG. This included examining changes in prediction accuracy for limited 2-channel FBAs based on bilateral channels of the bipolar montage. A FBA was trained and tested for each bilateral channel to examine the association between brain region and FBA accuracy. A Kruskal-Wallis test was used to determine if absolute PAD was different in groups related to training and testing location (9 bilateral groups: Fp1-F3/Fp2-F4, F3-C3/F4-C4, C3-P3/C4-P4, P3-O1/P4-O2, Fp1-F7/Fp2-F8, F7-T3/F8-T4, T3-T5/T4-T6, T5-O1/T6-O2, Fz-Cz/Cz-Pz). As a further adversarial test, we tested variations of channel laterality and channel location in 18 channel D1 data to gauge changes in FBA accuracy with respect to the spatial location of EEG channels. Potential differences in channel laterality were assessed by swapping left hemisphere EEG channels with right hemisphere EEG channels prior to training and testing. The same procedure was used to assess differences in anteroposterior directions, where anteriorly positioned (e.g. frontal) electrodes were swapped with posteriorly positioned electrodes (i.e. parietal, occipital) prior to training and testing. We also examined the temporal variation in FBA across all available epochs, i.e., from the start to the end of the recording period to observe optimal times to evaluate FBAs during N1 to N2 transitions.

We also performed additional sleep stage-specific tests to ascertain the performance accuracy of FBAs across wake, N2, N3 and REM sleep. Using Res-NN and GPR models, we estimated FBAs for all available EEG epochs for children in D2 (based on availability of sleep states). Age prediction within N2 sleep states were found to be the most accurate within our data (**Supplementary Table 3**).

#### Validation

We re-trained models using all available data in D1 and performed an out-of-sample validation on EEG data from D2. As D2 contained a limited number of recording electrodes, we used the same electrodes when training on D1 in a bipolar configuration (F4-C4 and C4-O2). Furthermore, only periods of N2 of D1 (10 minute segments) were included in the training dataset. We compared the wMAE between predicted and chronological age from the 10-fold cross-validation results from D1 to D2 to determine if the accuracy of the FBA trained on D1 was preserved when applied to D2. All validation analyses performed for the Res-NN model were repeated for the GPR model (see **Supplementary Figure 12** and **Supplementary Figure 14**).

To test site differences in EEG recordings (**Figure 5c**), we compared the distribution of EEG feature values using Kolmogorov-Smirnov tests corrected for multiple comparisons (Bonferroni’s method). We used a small random sampling of EEG features within several age defined bins (bin width was 1 year) to minimise the effect of age on feature distribution. Here, differences between EEG features per site and across age were denoted by the number of times the null hypothesis (EEG features between sites were from the same distribution) was rejected across 1000 samplings of each bin (*n* = 30 samples without replacement per feature per bin). The comparisons were also encoded to delineate if the distributional difference in EEG feature resulted in values greater than expected (older appearing) or less than expected (younger appearing) when comparing D1 to D2. At an acquisition level, we also estimated the total (summed) band power of the raw EEG signal, between 70 and 80 Hz, using Welch’s power density spectral estimate, to determine whether significant differences in the high frequency noise floor were present across sites (unpaired t-test).

#### Growth Charts

We generated growth charts for the FBA based on a limited 2-channel EEG using a combined D1 and D2 dataset. Centiles were estimated using generalised additive models with a protocol similar to the World Health Organization, WHO ^40^ child growth charts for height and weight for age. Here, we used the GAMLSS package ^41^ in RStudio (version 1.4.1717). These centiles were optimised by a Box-Cox-power exponential distribution with a cubic spline smoothing function, with distribution GAMLSS parameters *sigma* set as a cubic spline fit over age (df = 3) with *nu* = 1 and *tau* = 1. PAD was adjusted by significant factors uncovered during regression analysis (Eq 3.) to ensure that the FBA values presented in the growth chart were bias-free (i.e., accounting for confounding effects), as per best practices in brain age analyses ^3,8,42,43^. The “adjusted PAD” was used for subsequent group-based assessments alongside comparisons of centile-based values (which are inherently age-adjusted). Based on age corrected FBA values, centiles were estimated at the 3^rd^, 15^th^, 50^th^, 85^th^ and 97^th^ centiles as per WHO guidelines.

We also compared the performance of our FBA chart with paediatric head circumference and height measures. Using respective reference centiles across the paediatric ranges ^44,45^, we generated 1000 simulated training and testing cohorts from similar distributions of age, sex and cohort size to the combined D1 and D2 dataset. Using 10-fold cross-validation, we then trained and tested a GPR model with head circumference or height as input, and evaluated the results using MAE and wMAE.

Statistical tests were chosen on the basis of normality via a Lilliefors test^46^, where appropriate. If normality was met, group differences were examined by unpaired or paired t-tests.^47^ Evaluations of PAD and centiles to detect altered neurodevelopment as a ‘proof-of-concept’ were tested using t-tests, where the null hypothesis tested was that children with Trisomy 21 will have a PAD that is not different from children with typical neurodevelopment. Where appropriate, effect sizes in our study were reported using Cohen’s *d*. All statistical tests employed for analyses were two-sided and the level of significance was 0.05.

#### Ethics

The human research ethics committee at QIMR Berghofer Medical Research Institute approved the study (No. P3736, P3727). For D1, the Institutional Research Review Board at Helsinki and Uusimaa Hospital district approved the study (HUS/244/2021) including waiver of consent due to the retrospective collection of data acquired as part of standard of care. Ethics approval for the use of D2 and children with Trisomy 21 was granted by Children’s Health Queensland (LNR/2021/QCHQ/73595) including waiver of consent approved under the Public Health Act 2005 (PHA 73595) to analyse the retrospective cohorts.

## RESULTS

### Functional brain age

The FBA generated by the Res-NN was the most accurate predictor of chronological age (**Figure 3a-b**; MAE = 0.56 years, weighted MAE = 0.85 years, R^2^ = 0.96 and RMSE = 0.82 years; 10 fold cross-validation on D1, *n = 1056*), providing relatively uniform age predictions across the age range, with a median deviation of 10% from chronological age (see **Figure 4a**). Our comparative benchmarking of the Res-NN against a GPR approach and physical growth measures indicated that the neural network architecture exceeded (i) the age estimation accuracy of a GPR model (Cohen’s *d* = 0.31, *p* = 4.2 x 10^-23^, t-statistic = 10.1, paired t-test) which had a MAE of 0.79 years (**Supplementary Figure 12c, d**; wMAE = 1.06 years, R^2^ = 0.93 and RMSE = 1.09 years; see also **Supplementary Figure 12a, b** for comparison to Res-NN); (ii) the highest-performing individual EEG feature predictor (5^th^ percentile of EEG amplitude; MAE = 1.28 years, wMAE = 1.55 years, R^2^ = 0.82 and RMSE = 1.61 years, Cohen’s *d* = 0.59, *p* = 1.1 x 10^-69^, t-statistic = 19, paired t-test; **Supplementary Table 2**); iii) a prediction based on head circumference (estimated MAE = 1.72 years, wMAE = 2.54 years, Cohen’s *d* = 1.04, *p* =2.4 x 10^-143^, t-statistic = 27, unpaired t-test – see Methods for details); and iv) a prediction based on the mean age (MAE = 3.50 years, wMAE = 3.90 years, Cohen’s *d* = 1.3, *p* = 1.5 x 10^-266^, t-statistic = 40.5, paired t-test; n.b. this was performed to determine an upper bound for MAE based on the age distribution of the cohort).

### FBA model interpretability

To ascertain the presence of any bias in the data, we first examined the effects of age and sex on the FBA. The residual error between FBA and chronological age (referred to here as the predicted age difference or PAD) indicated a bias in FBA proportional to age (*β* = −0.036, *p =* 3.1 x 10^-5^, *n =* 1056, df = 1052). There were no significant differences in PAD between males and females (Cohen’s *d* = 0.04, *p =* 0.41, t-statistic = 0.83, unpaired t-test, *n =* 1056, df = 1054). Interactions between age and sex did not confound the relationship between PAD and age but a significant interaction with age was observed; PAD decreased with age (PAD ∼ sex: *β* = 0.036, *p* = 0.61; PAD ∼ age*sex: *β* = 0.009, *p* = 0.43, PAD ∼ age^2^: *β* =-0.011, *p* = 2.6 x 10^-14^, *n =* 1056, df = 1054).

We observed that early layers of network activations showed distinct age related clustering (**Figure 1c**; see also **Supplementary Figure 1** for general architectures) a pattern that becomes increasingly resolved with network depth (**Figure 4b**). We also observed that network activations were related to a range of EEG amplitude, frequency and entropy-based features. We found that EEG features, used in our GPR predictor, were highly correlated with activations present across early, middle and late stages of the NN architecture (median R^2^ = 0.50, IQR = 0.32, across all 32 features), suggesting that the Res-NN architecture captures several fundamental time and frequency domain characteristics of the EEG signal that are measureable with independent summary measures within its training phases (see **Supplementary Figure 8** and **Supplementary Figure 9**. Additionally, our GPR predictor derived from all 32 EEG features (R^2^ = 0.93, MAE = 0.79 years, wMAE = 1.06 years), consistently outperformed GPR predictors derived from a subset of features: only amplitude features (Feature IDs 1 to 7: R^2^ = 0.85, MAE = 1.10 years, wMAE = 1.30 years), only frequency features (Feature IDs 8 to 19: R^2^ = 0.90, MAE = 0.94 years, wMAE = 1.20 years) and only entropy-based EEG measures (Feature IDs 20 to 32: R^2^ = 0.91, MAE = 0.90 years, wMAE = 1.10 years). These relationships suggest that EEG features can be viewed as data surrogates that track the behaviour of neural network activations.

In addition to these tests, we also observed that EEG electrode location on the scalp and the timing of an epoch during an EEG segment (i.e., with respect to transitions between N1 and N2 sleep) influenced the FBA. FBA accuracy was significantly affected by the location of training electrode (*p* = 1.4 x 10^-31^, Kruskal-Wallis test). The accuracy of a 2-channel FBA was higher for posterior channels (**Figure 4c**), e.g. central, occipital (average MAE = 0.73 years), whereas anterior channels (e.g. frontal) had lower accuracies (average MAE = 0.83 years). Applying the predictor to data with swapped anterior and posterior channel positions resulted in a reduced performance accuracy (MAE = 1.11 years) whereas data with left hemisphere channels swapped with right hemisphere channels did not alter overall performance accuracy (MAE = 0.56 years). Temporally, the accuracy of the FBA was highest during the transition between N1 and N2 sleep states (**Figure 4d**; MAE = 0.61 years). Taken together, averaging FBA estimates across time and space improved overall accuracy.

### Validation

We then validated the FBA using a secondary dataset (D2) composed of 723 children recorded at the Queensland Children’s Hospital, Australia. To homogenise datasets across the different recording configurations in D1 and D2, we re-trained D1 on a 2-channel bipolar montage (F4-C4, C4-O2) and selected only periods of N2 sleep. In applying this retrained model to the validation set (trained D1 2-channel, tested D2 2-channel), the MAE of the FBA was significantly higher (**Figure 5a**; MAE = 2.17 years; wMAE = 2.27 years; R^2^ **=** 0.66; and RMSE = 2.05 years, Cohen’s *d* = 0.8, *p* = 2.4 x 10^-50^, t-statistic = 16.6, unpaired t-test) when compared to 2-channel models trained and tested on D1 and D2 individually (**Table 1** for Res-NN, **Supplementary Table 4** for GPR). Notably, the validation performance still outperformed the estimated accuracy of simulated head circumference based models (wMAE = 2.54 years).

**Table 1.** Overall performance of FBA across datasets. The performances of the FBA, using a Res-NN model, across all training, test, and combined datasets. R^2^, RMSE, MAE, and wMAE are shown. CI is confidence interval, ^a^only N2 from D1 used, ^b^10-fold cross-validation.

| Train | # channels,<br># recordings,<br># epochs | Test | # channels,<br># recordings,<br># epochs | $R^2$ | RMSE<br>(in years;<br>95% CI) | MAE<br>(in years; 95%<br>CI) | wMAE<br>(in years;<br>95% CI) |
| --- | --- | --- | --- | --- | --- | --- | --- |
| D1 | 19, 1056,<br>30624 | D1<sup>b</sup> | 19, 1056,<br>30624 | 0.96 | 0.82<br>(0.78 - 0.90) | 0.56<br>(0.52 - 0.59) | 0.85<br>(0.69 - 1.02) |
| D1<sup>a</sup> | 2, 1056,<br>20064 | D1<sup>a,b</sup> | 2, 1056,<br>20064 | 0.93 | 1.10<br>(1.08 - 1.26) | 0.77<br>(0.72 - 0.82) | 1.22<br>(0.96 - 1.48) |
| D1<sup>a</sup> | 2, 1056,<br>20064 | D2 | 2, 723,<br>13737 | 0.66 | 2.76<br>(2.61 - 2.89) | 2.18<br>(2.05 - 2.29) | 2.27<br>(1.90 - 2.65) |
| D2 | 2, 723,<br>13737 | D2<sup>b</sup> | 2, 723,<br>13737 | 0.78 | 1.53<br>(1.72 - 1.91) | 1.45<br>(1.37 - 1.53) | 1.66<br>(1.37 - 1.96) |
| D1<sup>a</sup>+D2 | 2, 1779,<br>33801 | D1<sup>a</sup>+D2<sup>b</sup> | 2, 1779,<br>33801 | 0.88 | 1.41<br>(1.49 - 1.64) | 1.09<br>(1.04 - 1.14) | 1.51<br>(1.30 - 1.73) |

The decrease in FBA performance was attributed to three key site differences: age distribution, number of EEG channels, and site-specific differences of the EEG across age. The effect of age distribution between sites (**Table 1**) contributed accounted for an approximate net increase of 0.24 years in the MAE, when comparing MAE to wMAE from the primary training data (D1 cross-validation) versus external validation data (D2 cross-validation). Similarly, a reduction in the number of EEG channels from 18 to 2 resulted in an increase of 0.37 years in the wMAE (**Table 1**). The effectiveness of the Res-NN in capturing fundamental characteristics of the EEG signal (**Figure 5b**), corresponded well to age-specific differences in individual summary EEG features were observed across sites (**Figure 5c** and **d**). Approximately 38% of EEG features/age bins combinations differed significantly across age and sites with 194 out of 512 hypothesis tests meeting significance at p<0.05 following correction for multiple comparisons. Additionally, we found noteworthy distinctions in spectral estimates of the EEG recording noise floor between sites, whereby EEG recordings in D1 exhibited a higher noise floor compared to D2 (70 to 80 Hz band power; Cohen’s *d* = 0.37, *p* = 1.7 x 10^-12^, t-statistic = 7.1, unpaired t-test). Training a FBA model constructed from a combined dataset D1 and D2 considerably improved overall prediction accuracy with a MAE of 1.04 years for D2 (*n* = 723) suggesting site specific differences were incorporated into the model. **Supplementary Table 5** summarises model performances across test folds for all datasets, respectively.

### Functional growth charts

We next combined both datasets (D1 + D2) to generate a FBA ‘growth chart’, wherein generalised additive models were applied to construct age-appropriate centiles ^40,48^. The resultant FBA had an MAE of 1.09 years with a wMAE of 1.51 years, an R^2^ of 0.88 and an RMSE of 1.41 years (**Figure 6a**; see also **Table 1**). Further, FBA growth charts based on an age-stratification of infants (0 to 2 years; MAE = 0.40 years) and children (2 to 18 years; MAE = 1.34 years) indicated a high degree of accuracy relative to their respective age group (**Supplementary Figure 13**).

**Figure 6.**
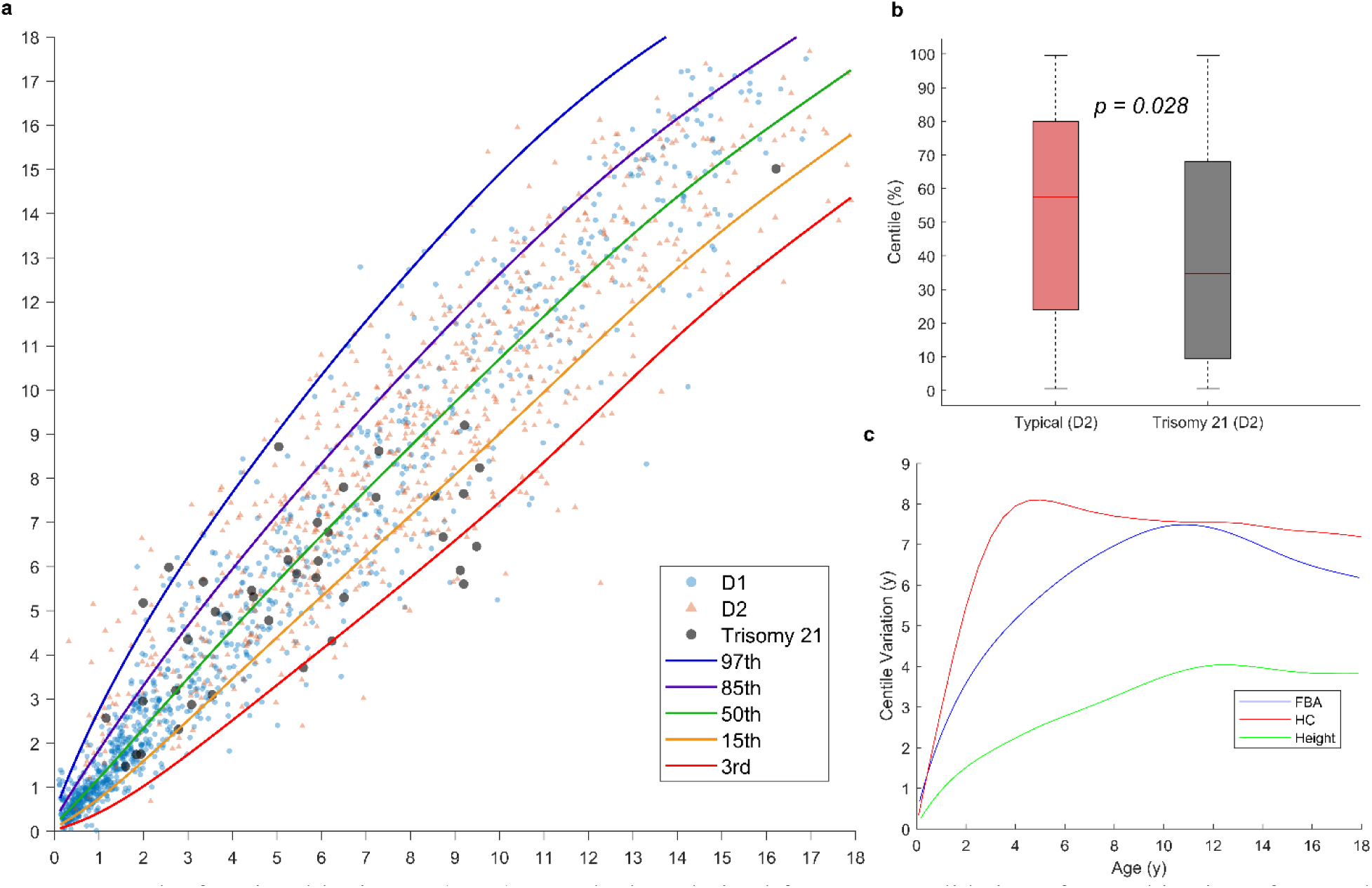
The functional brain age (FBA) growth chart derived from cross-validation of a combination of D1 and D2 datasets. **a.** The chart is based on the FBA derived from the Res-NN model. Translucent colored triangles represent individual EEG recordings from D1 (blue) and D2 (red) datasets. The 3^rd^ (red), 15^th^ (yellow), 50^th^ (green), 85^th^ (purple) and 97^th^ (blue) centiles are indicated. Children with Trisomy 21 (black dots) have been plotted alongside D1 (blue) and D2 (pink) children. **b.** Differences in relative PAD between children with typically developing neurodevelopment and children with Trisomy 21. Significance values (*p <* 0.05) were determined by conducting an unpaired t-test between groups, where all data was checked for normality. Typically developing groups (*n* = 723; blue) and Trisomy groups (*n* = 40; pink) from D2 are plotted as violin plots, with the median (black line) and interquartile ranges (rectangles) shown, **c.** Variation in paediatric predictors of age. The variation (in years) represents the difference between the 97^th^ and 3^rd^ centiles across age for the FBA (in blue), head circumference (HC, in red) and height (in green). HC and Height predictors were simulated with the same age distribution as the combined D1 and D2 dataset.

The practical utility of an FBA is that it enables stratification of children, by quantifying brain functions associated with a child’s diagnostic status and underlying neurodevelopmental issues. To demonstrate this, we compared typically developing children from D2 with an additional small cohort of children from the same site whom were diagnosed with Trisomy 21 (*n* = 40; 29/40 were recorded at less than 7 years of age). The PAD was significantly lower in children with Trisomy 21, despite having normal sleep studies, than typically developing children in D2 (PAD adjusted for age effect: *p* = 5.3 x 10^-4^, t-statistic = 3.5, unpaired t-test, n = 763, df = 761; centile-based: Cohen’s *d* = 0.36, *p* = 0.028, t-statistic = 3.5, unpaired t-test, n = 763, df = 761; **Figure 6b**). This finding of significantly lower PAD was consistent across other combinations of cohorts: (i) typically developing children from D1 only versus children with Trisomy 21 (*p* = 8.7 x 10^-3^, t-statistic = 2.6, unpaired t-test, df = 1094) and (ii) typically developing children from D1 + D2 versus children with Trisomy 21 (*p* = 8.4 x 10^-3^, t-statistic = 2.6, unpaired t-test, df = 1817). No significant differences in age and sex were found in children with Trisomy 21 (*p* = 0.61, t-statistic = 0.5, unpaired t-test). **Supplementary Table 6** summarises all further comparisons including effect sizes and sex differences. The observations suggest that at the group level, deviant neurodevelopmental trajectories in children with Trisomy 21 translate to delayed maturation of their cortical function.

Finally, we benchmarked our FBA model against conventional growth chart trajectories of head circumference and height in children.^44,45^ Here, the maximal variation of age for the FBA (difference between 3^rd^ and 97^th^ centiles) falls between head circumference and height for age for a simulated cohort with similar age and sex demographics to D1 and D2 combined (**Figure 6c**). This indicates that the variation in FBA, for a typically developing cohort, as per our estimated centiles, are relatively smaller for younger children in comparison to larger variations for children above 10 years of age; a trajectory that is generally observable in both charts based on anthropometric and neuroimaging measures across the lifespan.^5,6,44,45^ This variation is also likely attributed to the distribution of age presented in D1 + D2, where samples of adolescents only account for 20% of the combined dataset. Our FBA growth chart thus exhibits comparable age variability to that of widely-used physical growth charts. Code for converting EEG into FBA and centiles are available (details in *Data sharing statement*).

## DISCUSSION

In infants, children and adolescents, EEG activity exhibits clear, consistent, and rapid changes with age.^13,49,50,51^ We formalise this knowledge using a targeted deployment of deep learning algorithms to form a prediction of EEG age (FBA) underpinned by key human operator expertise and decision-making in specific stages of the process. We achieved an accurate FBA prediction when applying deep neural networks directly to the EEG signal, relying on a summary of only brief epochs (60 seconds) within a 10 to 15 minute segment recorded during N1/N2 sleep providing similar accuracies with widely used anatomical growth charts.^40,44,45^ The proposed FBA demonstrated state-of-the-art age prediction accuracy, was validated in an independent cohort and detected group level maturational delays in a small cohort of young children with a defined neurodevelopmental disorder.

Our FBA estimates had MAEs comparable to the highest performing MRI-based^2,3,6,7,10,23^ and EEG-based studies^24,25,26,27,28^ with reported MAEs in the literature ranging from 1.0 to 4.6 years compared to our best wMAE of 0.88 years. The accuracy of the FBA may be directly attributable to the use of residual neural network architectures over conventional multivariable age regression approaches typically used in brain age studies. While individual features have significant correlations with age (**Supplementary Table 2**), the combination of these features provided a superior prediction of age. Training deep neural networks improved these predictions further, although the exact mechanism of this improvement is not entirely clear. We show that deep neural networks capture well-established EEG characteristics (such as amplitude, frequency, bursting behaviour, and entropy) by comparing features to internal network layer outputs and that the representation of these latent patterns improve FBA estimates through higher-dimensional abstractions of the EEG signal. By showing that individual EEG features correlate with the outputs of network layers, we highlight a demonstrable feature of the FBA during training. However, methods that attempt to explain the function of neural networks must be made with caution.^52^

The present FBA measures were developed solely using large EEG datasets that are routinely collected and widely available in hospitals worldwide. Although the performance error of the FBA increases with age, the relative accuracy of the FBA is comparatively uniform across the age range. Here, additional variations to FBA accuracy were linked to the spatial and temporal organisation of the EEG. The effect of spatial organisation for instance was primarily a frontal-occipital gradient which is a well-established phenomenon in the maturing EEG within this age group.^53,54^ Temporally, the accuracy of the FBA was maximum at the onset of N2 sleep characterised by the presence of sleep spindles which are key cortical signatures that emerge in the first few months of life and remain present in the EEG through adulthood.^31^ Improved EEG stability near the sleep state transition may involve capitalising on the absence of critical slowing within EEG dynamics at the beginning of these state-based transitions.^55^

The only manual selection done prior to our computational analyses was the identification of the first sleep spindle as a sign of N2 sleep, which was necessary to harmonise vigilance states across a cohort with a wide age range. There are several reasons as to why the N2 sleep state offers a well standardised vigilance state that can be considered much more homogeneous across individuals than compared to wake or other sleep states.^15^ The identification of N1/N2 states, which is marked in the EEG by the emergence of increased delta, sleep spindles, vertex waves and K-complexes, particularly in N2 sleep, are well studied, reliable EEG signatures^56,57^ across preclinical and clinical literature. A brief period of N2 sleep is also often recorded in routine EEG studies as it is rich in EEG signatures and known to be sensitive for observing pathological phenomena, (such as epileptiform events^58,59^), and is also minimally contaminated by the common artefacts due to movements. The EEG during N2 is also a well-known target for tracking neurodevelopment, with an initial increase in EEG amplitude during infancy followed by a steady decline into adolescence. Global spectral power shows a decrease in delta frequencies offset by a steady increase in relative alpha and beta band power with age (as observed in Feature IDs 13 to 17 versus age in **Supplementary Figure 10**), likely reflecting increasing dominance of spindle activity in EEG spectra with age.^56,60,61,62^

An unresolved question in this work is whether an FBA measured within other diverse vigilance states (e.g. resting, task, or other sleep states) could effectively enhance the accuracy of individualised assessment. Our additional tests during sleep and wake states (**Supplementary Table 3**) demonstrate the applicability of an FBA in these potential contexts. Obtaining consistent awake EEG in older cooperative children is feasible, but collecting several minutes of good quality EEG signals from alert infants and toddlers is difficult. Careful consideration is essential in harmonising of spontaneous EEG data, especially given the neurophysiological and behavioural variability during childhood.^63,64^ To enhance the signal-to-noise ratio in comparisons between younger and older children, it becomes crucial to ensure a larger pool of available EEG data for the younger age group.^64^ Defining normative variability margins in typical development via large consortia EEG datasets, such as the Healthy Brain Network (>3000 children^65^) and comparable hospital-based clinical EEGs^66^, are likely to provide clues into the scope of the FBA beyond the paradigm of N2 sleep.

The present study has some potential limitations. The performance of external validation was markedly lower than the overall performance of cross-validated results in each site independently. The drop-off in accuracy is due to several factors, namely: site specific differences, a lower electrode density, and inherent differences in acquisition of the EEG recordings. We showed that increasing the diversity of training data, by combining data from D1 and D2, mitigates this issue. However, to enhance accuracy, external validation datasets with diversity in geography, ethnicity, and socioeconomic status will improve the generalizability of the FBA. Despite the trade-off in performance accuracy, our externally validated results still outperformed measures such as head circumference simulated over the same age range.

Another limitation of the study is that all children included in this study were not representative of the larger, healthy paediatric population but rather a subgroup of children clinically referred from the primary care level to a tertiary care center for diagnostic assessment. In neurotypically developing populations, it is expected that around 5% may conceal potential subclinical pathologies^67^ – a trait notably observed among individuals falling outside the 3rd and 97th centiles on our growth chart (**Figure 6a**). Estimates of FBA in such groups are clinically interesting; however, it is essential to benchmark the FBA in healthy neurotypical cohorts, including the use of longitudinal data, to ensure further clarity and confidence in applying FBA to a broader paediatric population. The progressive refinement of FBA methods in neurotypical EEGs can enhance our understanding of how FBA models should navigate the balance between aleatoric uncertainty and epistemic uncertainty encountered in large datasets.

The FBA also appears to compensate for, or is indifferent to, growth spurts, hormonal and pubertal changes in both sexes, and other alterations to brain structure such as increased rates of cortical thinning in males during adolescence.^68,69^ This does not discount the fact that factors such as sex related differences in cortical activity exist across age, rather, that sex specific effects in the EEG were accounted for and adjusted out by the model.^50^ Future applications of the FBA could be used to study genuine sex-related differences in cortical maturation ^50^ by separating data according to biological sex at the training stage. This ability of trained models to inherently adjust for potential confounders is a key aspect of artificial intelligence (AI) methods in medicine and means that what we know about EEG and age should be reevaluated singularly in the context of the AI outputs. Growth charts can also be calibrated for diagnosis, prognosis, and stratification; here the optimal tradeoff between cohort heterogeneity, EEG acquisition, training data size and MAE is not entirely resolved, with evidence from MRI-based studies suggesting that the clinical utility is not necessarily inversely proportional to MAE.^6,70^

We propose the FBA as a measure that enables assessment of neurodevelopmental trajectories from infancy to adolescence. Rather than replacing or challenging existing techniques, the EEG-derived FBA is perhaps best seen as a valuable complement to support current modalities of neurodevelopmental assessment, offering a tool towards personalization that both benefits the patient and healthcare practitioner alike. While recognizing the FBA’s clinical potential, a series of targeted evaluations of the FBA within clinical populations are necessary to determine its efficacy prior to endorsing its widespread use. These extra studies are required not only to determine the clinical utility of the algorithm but to also enable other researchers and institutions to identify appropriate safeguards for decision safety and efficacy. We, therefore, publicly release the FBA prediction algorithm as an ‘online’ resource that facilitates the continual refinement of targeted algorithms for tracking childhood brain function and neurodevelopment.^71^

## Contributors

Conceptualization: K.K.I, J.A.R, M.W, S.V, N.J.S

Methodology: K.K.I, M.W, L.L, S.V, N.J.S

Investigation: K.K.I, S.V, N.J.S

Visualization: K.K.I, N.J.S

Funding acquisition: M.W, S.V, N.J.S

Project administration: K.K.I, L.L, S.V, N.J.S

Supervision: L.L, S.V, N.J.S

Writing – original draft: K.K.I, J.A.R, M.W, L.L, S.V, N.J.S

Writing – review & editing: K.K.I, J.A.R, M.W, J.C, S.J.V, A.K, L.M.H, L.L, S.V, N.J.S K.K.I, N.J.S, L.L, S.V accessed and verified the underlying data.

All authors read and approved the final version of the manuscript.

## Data sharing statement

Trained Res-NN and GPR models for age prediction are available on our GitHub repository, accessible at: https://github.com/brain-modelling-group/functional-brain-age. Data used to train and validate the FBA will be made available to researchers on reasonable request to S.V., N.J.S. K.I.: data sharing is subject to a material transfer agreement, approved by the legal departments of the requesting researcher and by all legal departments of the institutions that provided data and ethical clearances for the study. EEG feature matrices (z-scored) and child ages for each EEG recording in our training dataset (D1, 18 channel and 2 channel versions) and external validation set (D2, 2 channel version) are also available in our GitHub repository. These files have been pseudonymised. MATLAB and Python code for the prediction of functional brain age by Res-NN based methods on 18- and 2-channel EEG are available on our GitHub repository.

## Acknowledgements

We acknowledge our funding partners who supported this work, including: NHMRC Grant no. 2002135, The HUS Children’s Hospital/HUS diagnostic center research funds, Finnish Academy (335788, 332017), Finnish Paediatric Foundation (Lastentautiensäätiö) and Sigrid Juselius Foundation.

We would like to acknowledge and thank Dr Paula Sanz-Leon for technical assistance with the deployment of functional brain age software provided on our GitHub repository. For Dataset 1, we thank Mr Jarmo Mäki and other EEG technicians within the HUS diagnostic center in annotating EEGs, as well as Mr Eero Ahtola for assisting with the preprocessing of EEG data. For Dataset 2, we thank all technicians, nurses and doctors involved with data collection, overnight annotations and clinical assessment of polysomnography recordings conducted at the Respiratory and Sleep Medicine Department located within the Queensland Children’s hospital.

---

**Supplementary Figure 1.**
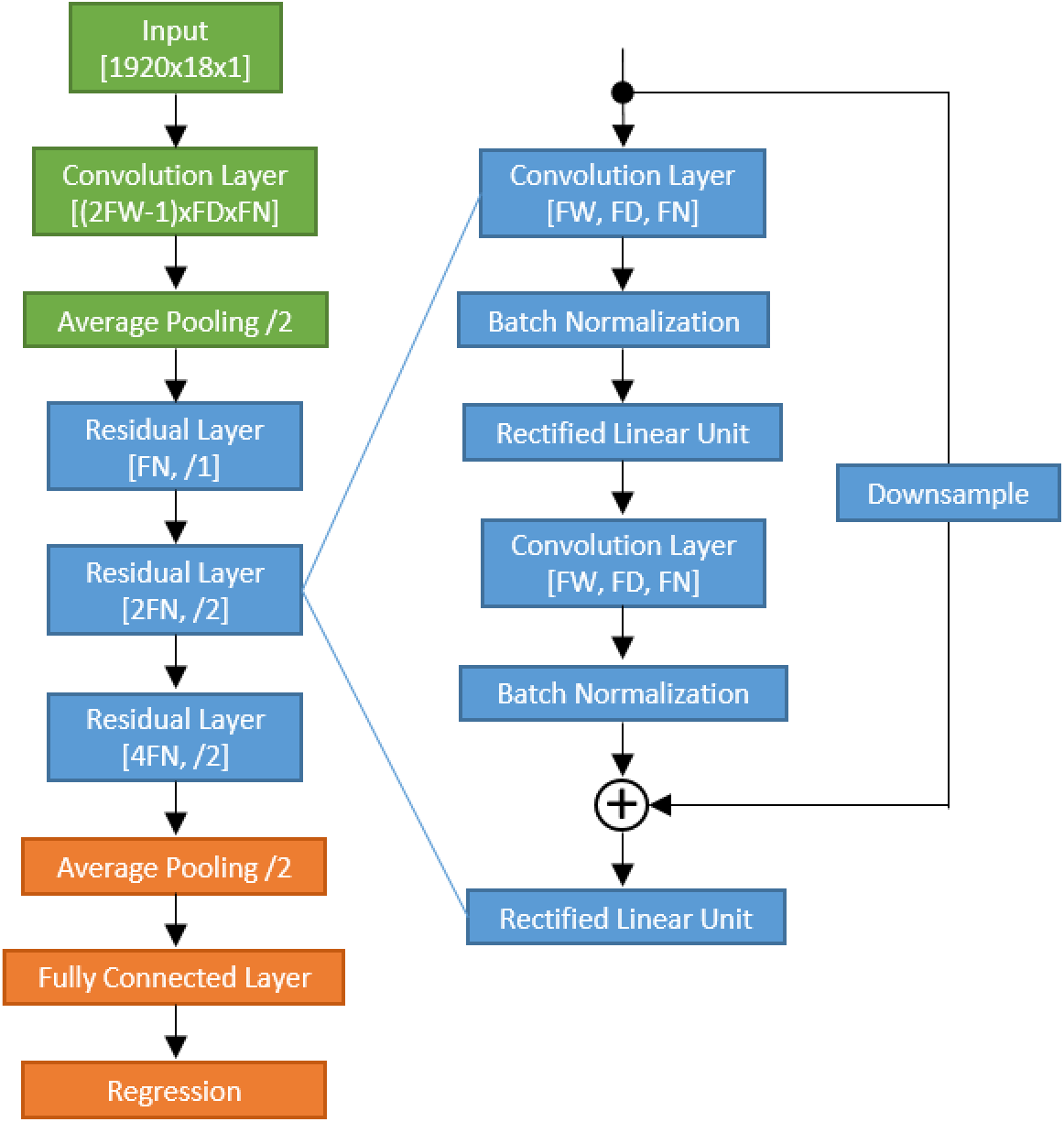
The general architecture of the residual neural network used in this work. For inception networks, we used elements of the inceptionv3 network in MATLAB with a final regression layer. The scale of this pre-trained network was too large to accurately train on the dataset used in our work, so we tested architectural aspects of the network rather than the entire network. Similarly to the residual neural network we added variability by changing the temporal filter width (FW), filter channel depth (FD) and filter number (FN) within the convolutional layers. We used the file generate_networks_v2.m to generate networks with different configurations and architectures (see the GitHub page for more details).

**Supplementary Figure 2.**
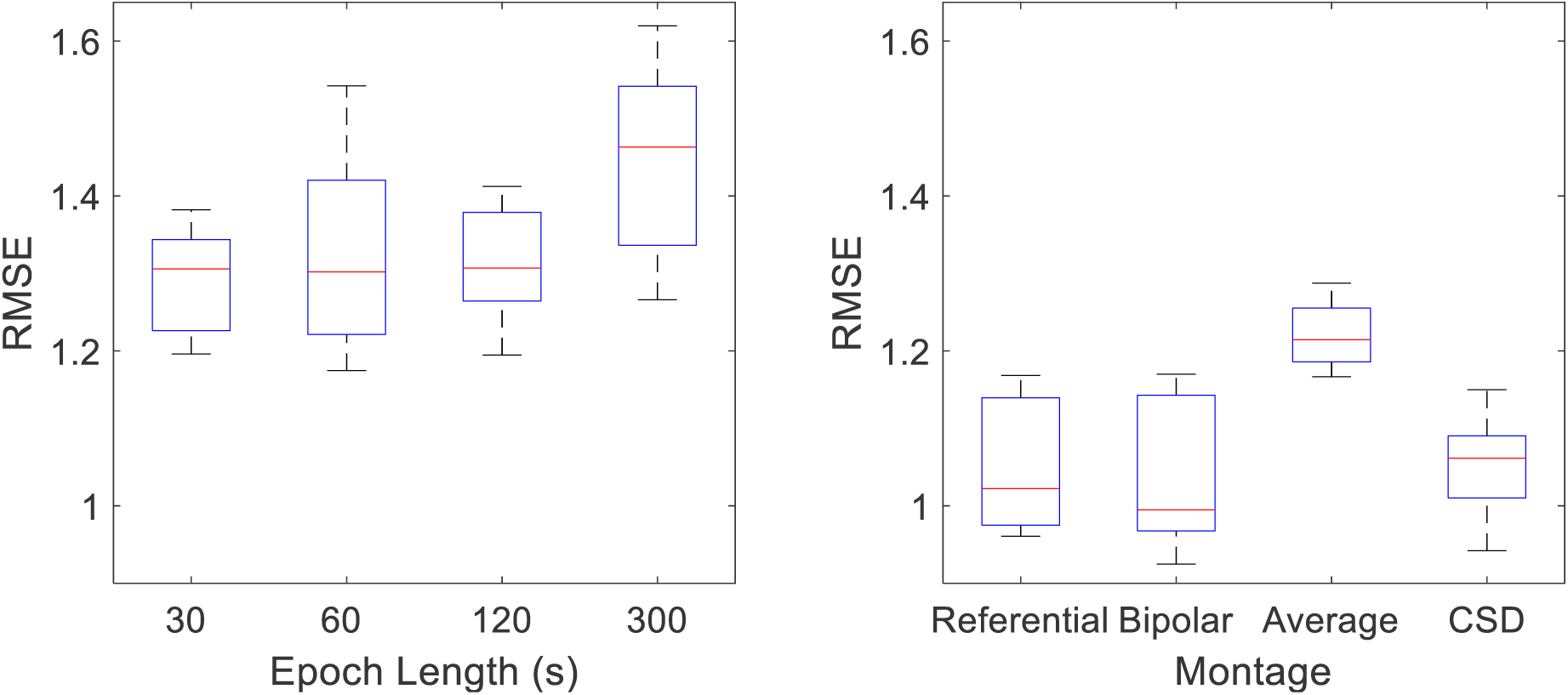
The effect of epoch length and montage on cross-validated RMSE on a random selection of 7 network architectures. An epoch length of 60s and the bipolar montage were selected *a priori*. Different EEG epoch durations (30 s, 60 s, 120 s, 300 s) and EEG montages (referential, bipolar, average, current source density) were tested. The root mean square error between predicted age (FBA) and age across all testing data from a 10-fold cross-validation was used to determine the optimal selection, with a minimum RMSE indicating the optimal results.

**Supplementary Figure 3.**
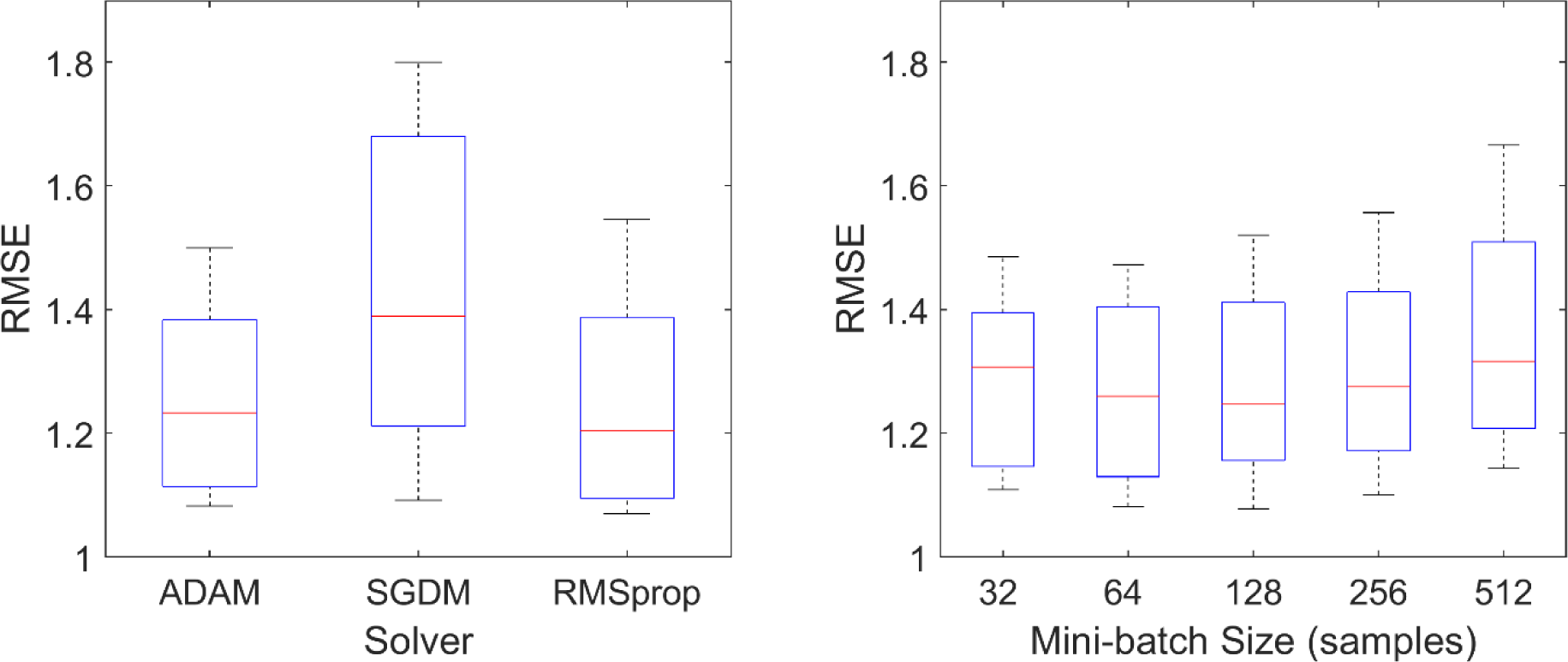
The effect of solver and mini-batch size on cross-validated RMSE on a random selection of network architectures, epoch length, and montage (*n* = 8 per group for solver and *n* = 7 per group for mini-batch size). For training the neural networks, two hyper-parameters of training were selected based on an initial run on dataset D1 (Solver Type and Mini-batch size). A random selection of network architectures were made and the network was trained and tested within a 10-fold cross-validation with the RMSE on the accumulated left-out test data used to select the hyper-parameters. The ADAM solver and a mini-batch size of 128 samples were selected.

**Supplementary Figure 4.**
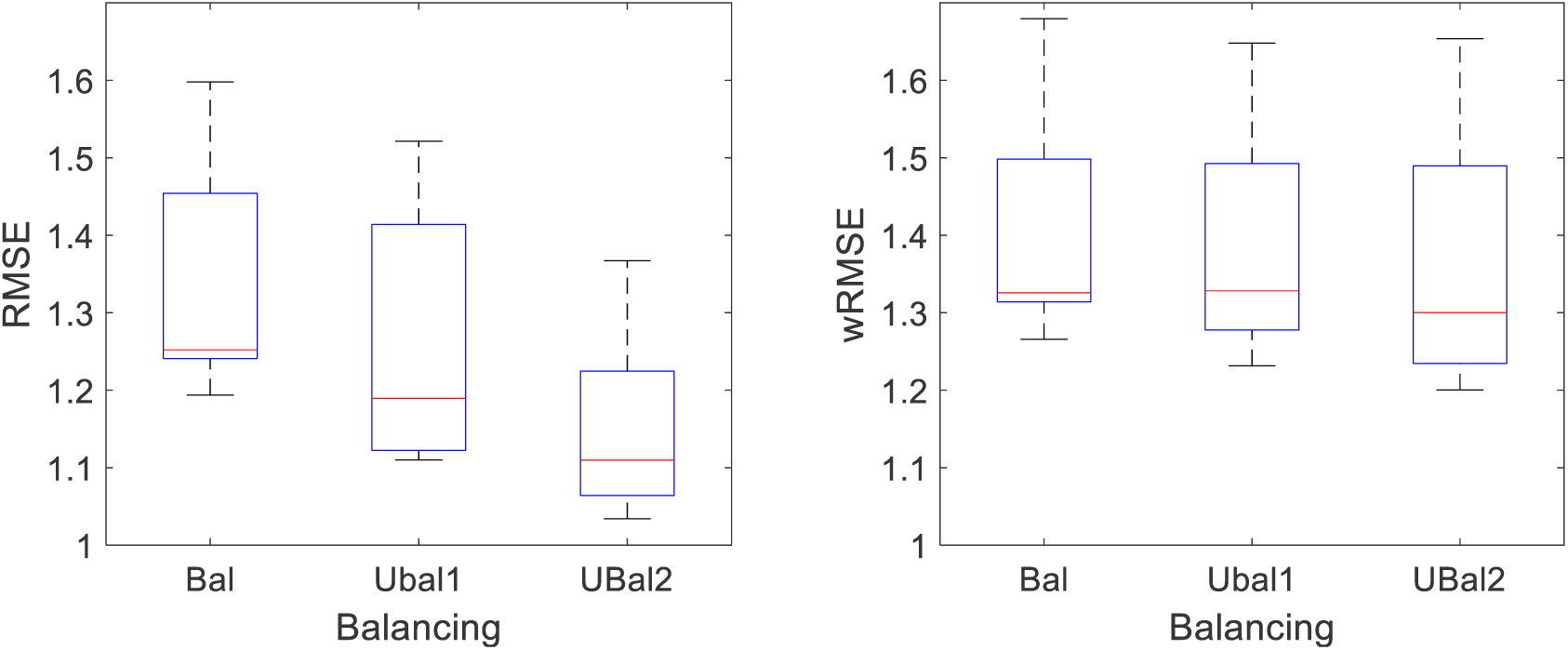
The general effects of data balancing on cross-validated RMSE for 18-channel data (D1 datasets). The left plot is the RMSE and the right plot is the wRMSE which is an RMSE averaged across age bins (yearly) rather than per EEG recording. In both cases, the increased diversity in recordings associated with unbalanced training dataset offers lower errors, although the reduction in error is smaller when considering wRMSE.

**Supplementary Figure 5.**
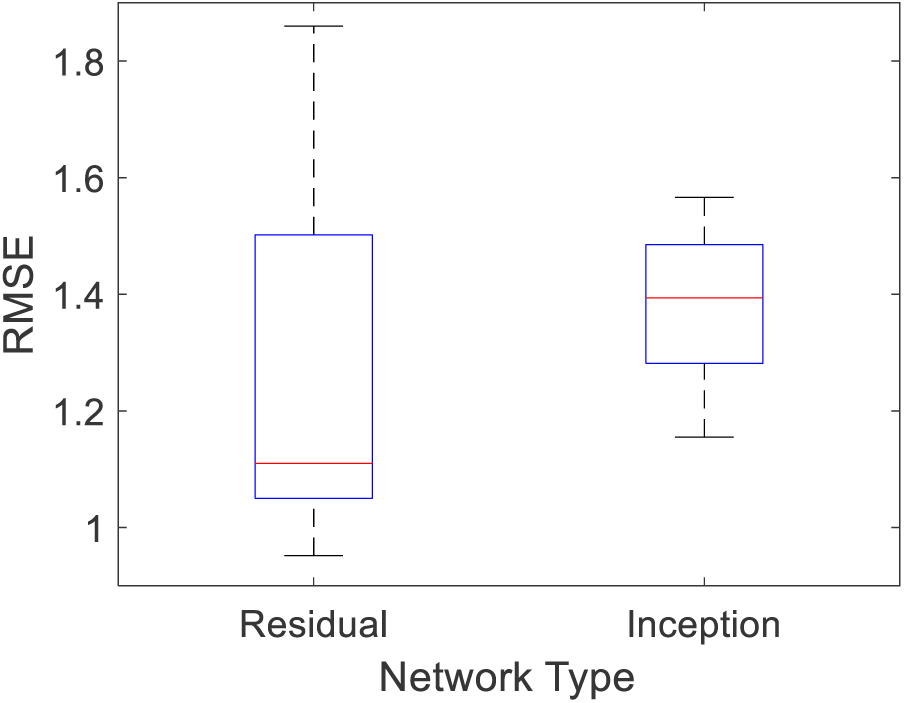
The general effects of data network architecture type (Residual vs Inception) on cross-validated RMSE for 18-channel data (D1 datasets). Once general EEG parameters and network training hyper-parameters were initialized, we tested two general architectures based on residual neural networks and inception based neural networks. We randomly selected 10 networks based on both architectures and computed RMSE. Based on these parameter evaluations, residual networks were selected as the most suitable candidate architecture for estimating age.

**Supplementary Figure 6.**
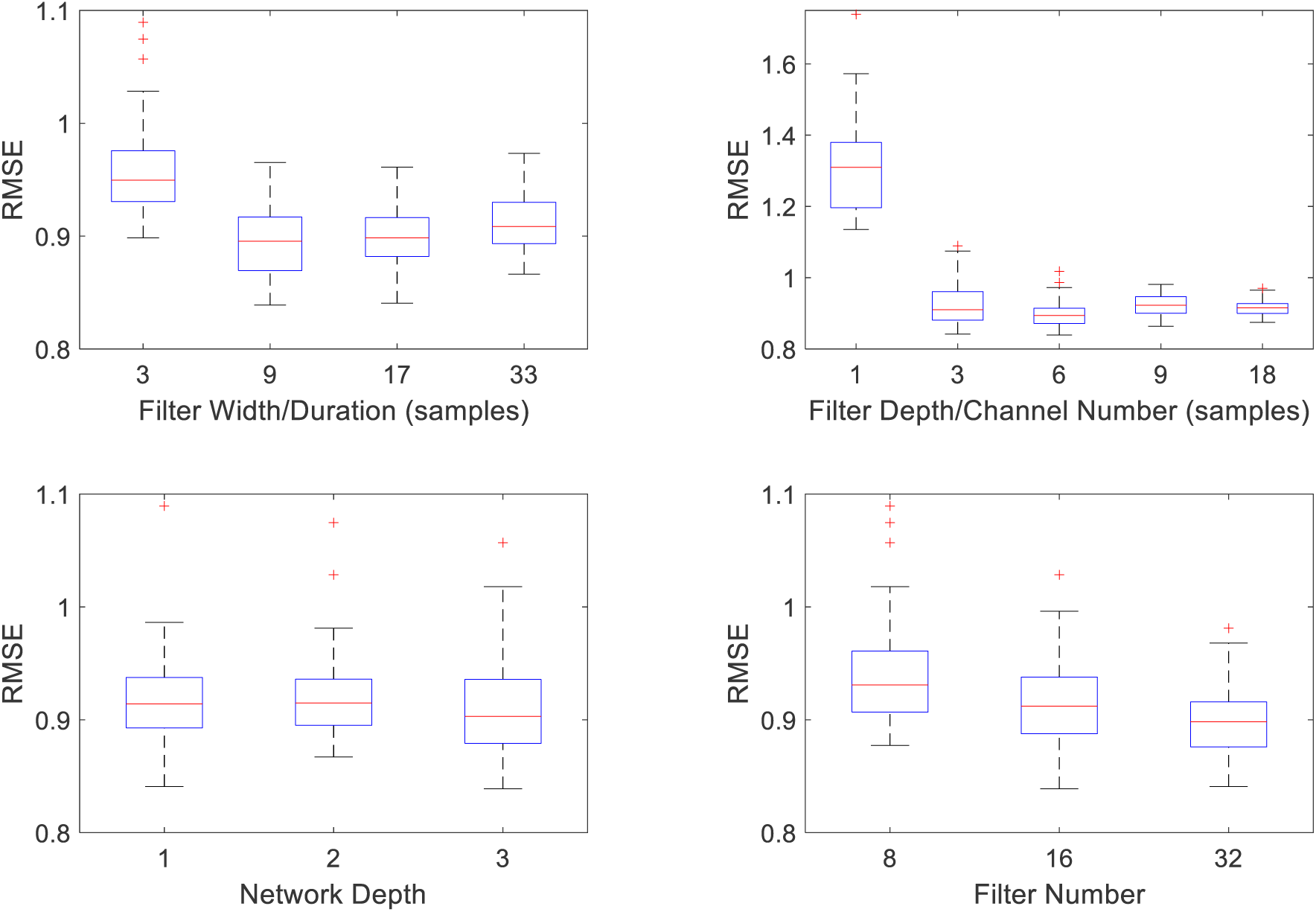
The general effects of residual network parameters on cross-validated RMSE for 18-channel data (D1 dataset). Internal single fold cross-validation averaged across all 10 training folds selected. Note that, as filter depth of 1 channel had a considerably higher RMSE than other filter depths, RMSE points associated with a 1 channel network were removed before generating the remaining boxplots for visualization purposes.

**Supplementary Figure 7.**
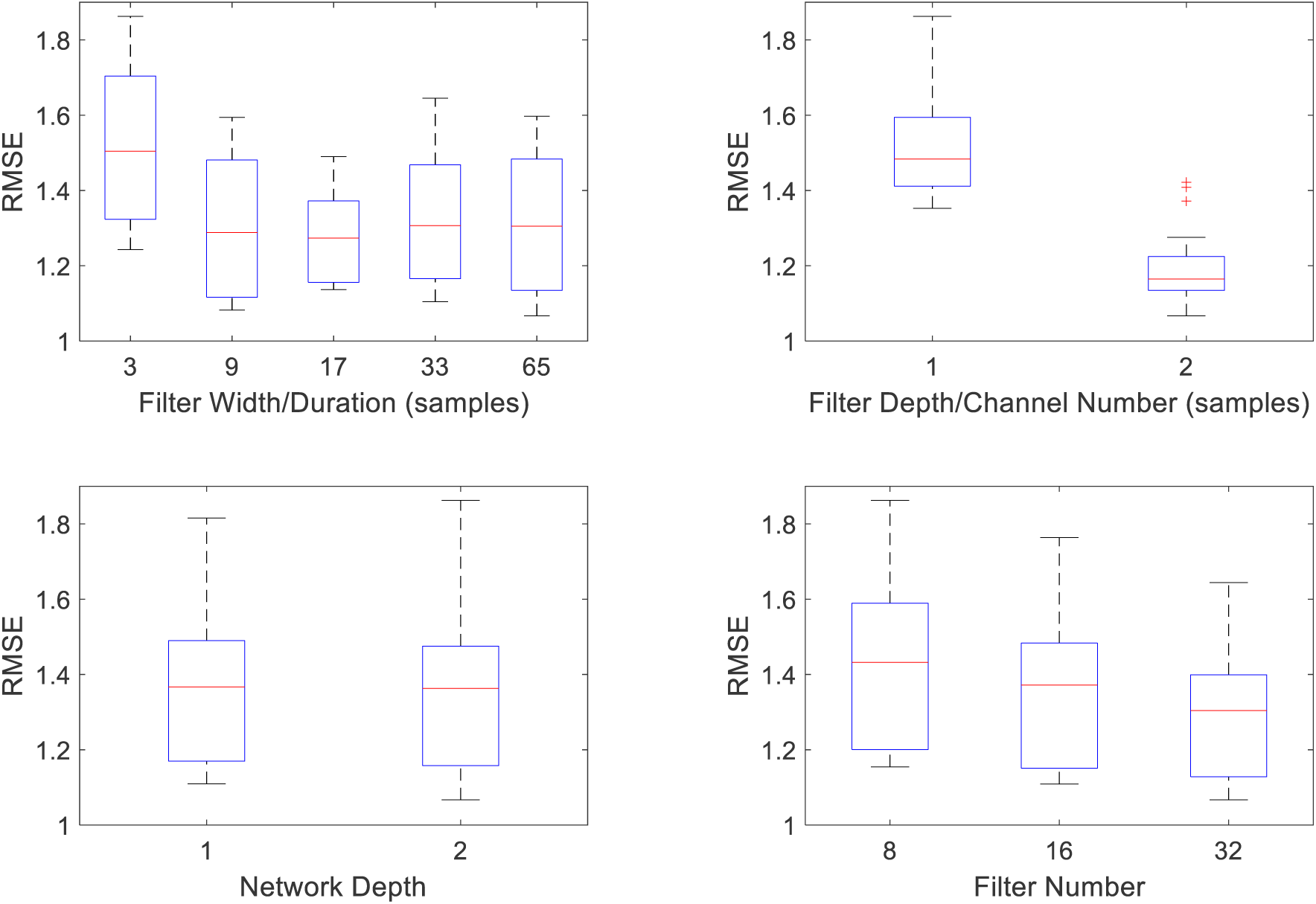
The general effects of residual network parameters on cross-validated RMSE for a 2-channel FBA based on dataset D1. This resulted in 60 different network architectures. The optimal combination of network parameters selected were a filter width of 9 samples (*n* = 18 per group), filter depth of 2 channels (*n* = 30 per group), network depth of 2 (*n* = 30 per group), and a filter number of 2 (*n* = 20 per group): a total of 906,657 learnable parameters.

**Supplementary Figure 8.**
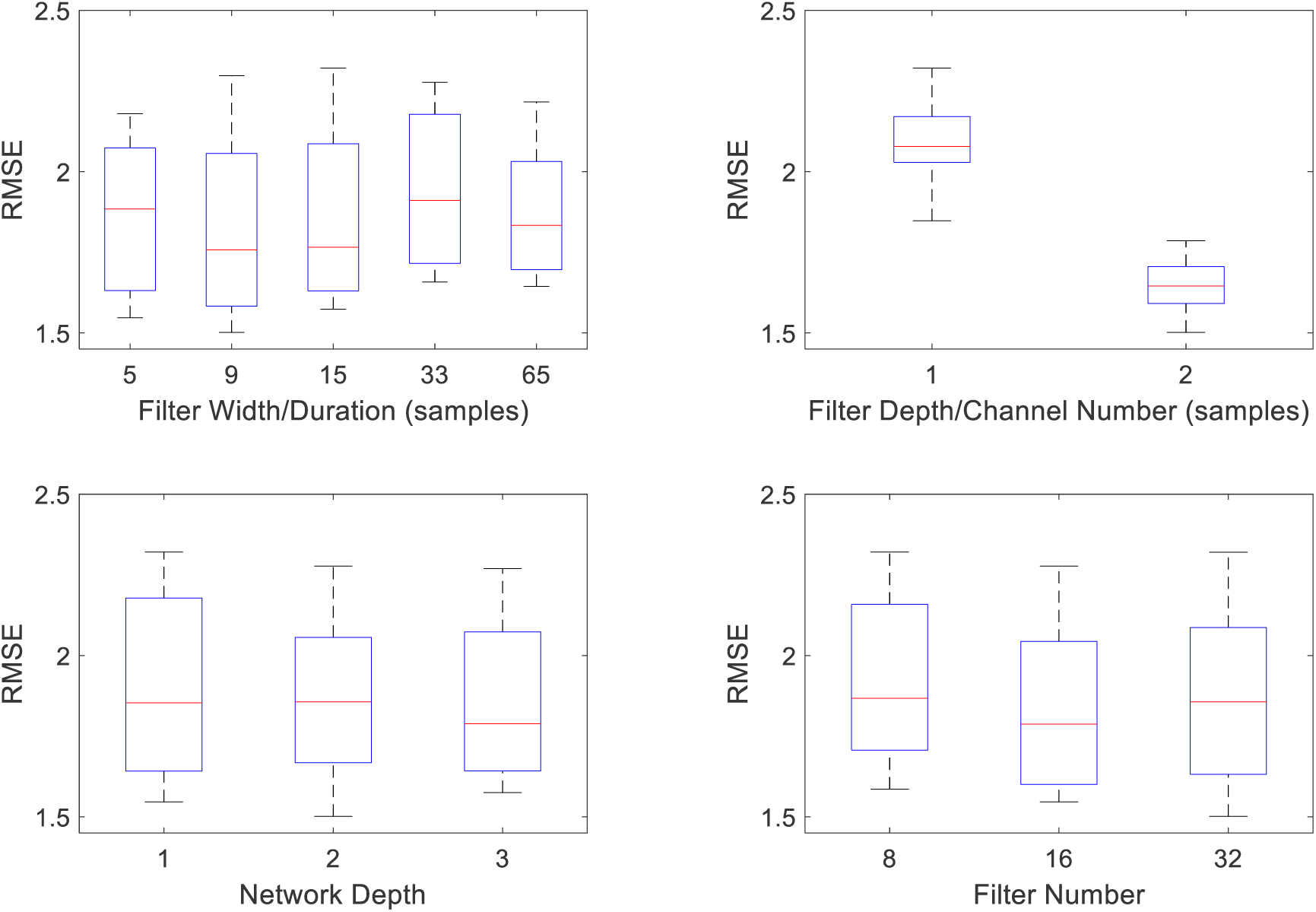
The general effects of residual network parameters on cross-validated RMSE for 2-channel data (D2 dataset). This resulted in 90 different network architectures. The optimal parameters were a filter width of 9 samples (*n* = 18 per group), filter depth of 2 samples (*n* = 45), network depth of 2 (*n* = 30 per group), and a filter number of 32 (*n* = 30 per group): a total of 906,657 learnable parameters.

**Supplementary Figure 9.**
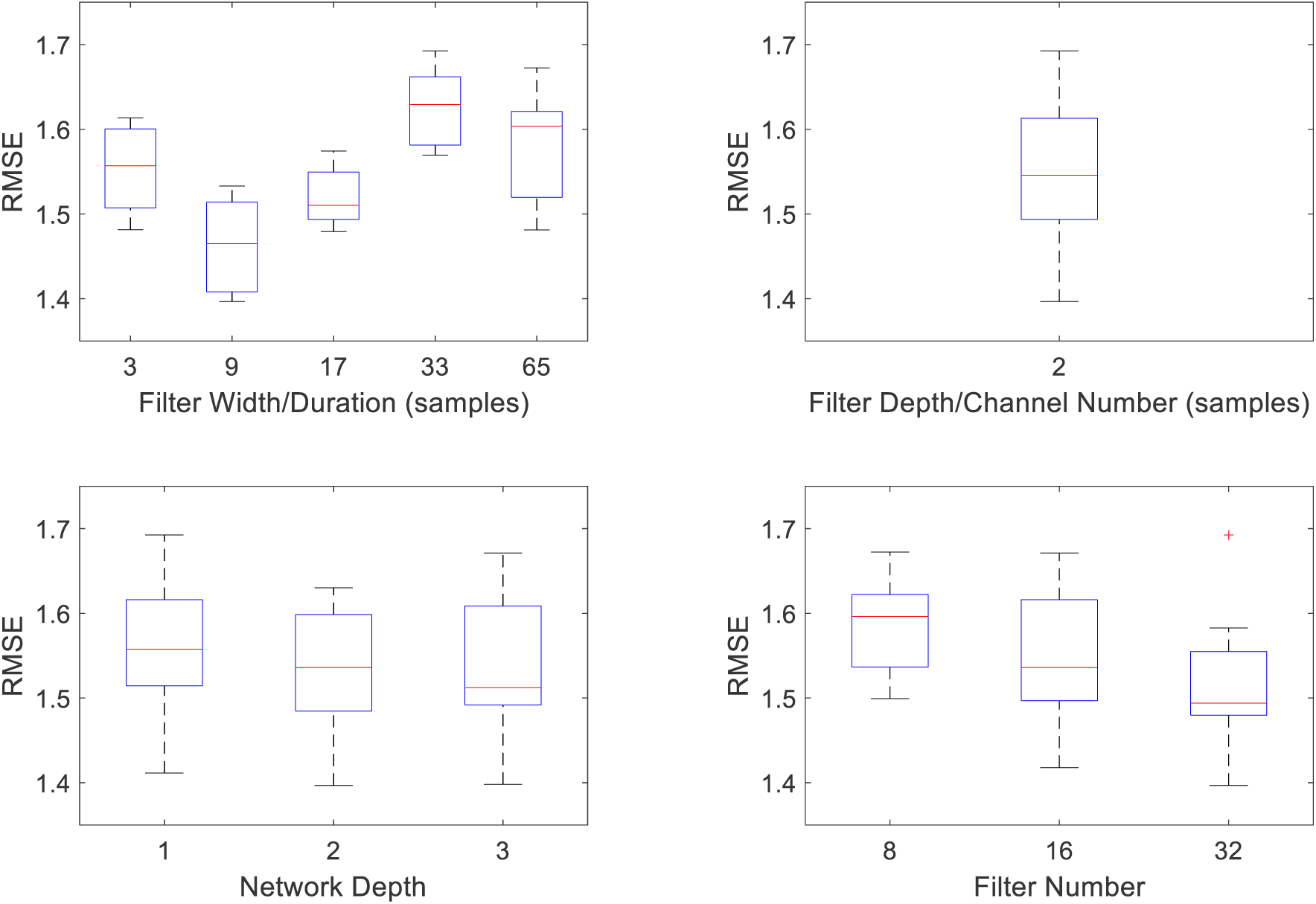
The general effects of network parameters on cross-validated RMSE for 2-channel data (D1 and D2 combined). This resulted in 45 different network architectures. The RMSE of an internal single fold cross-validation, average across all 10 training folds selected a filter width of 9 samples (*n* = 9 per group), a network depth of 2 (*n* = 45 per group), network depth of 2 (*n* = 15 per group), and a filter number of 32 (*n* = 15 per group): a total of 906,657 learnable parameters. Note, we did not vary the filter depth/channel number as previous experiments on D1 and D2 alone showed that 2 channels was optimal.

**Supplementary Figure 10.**
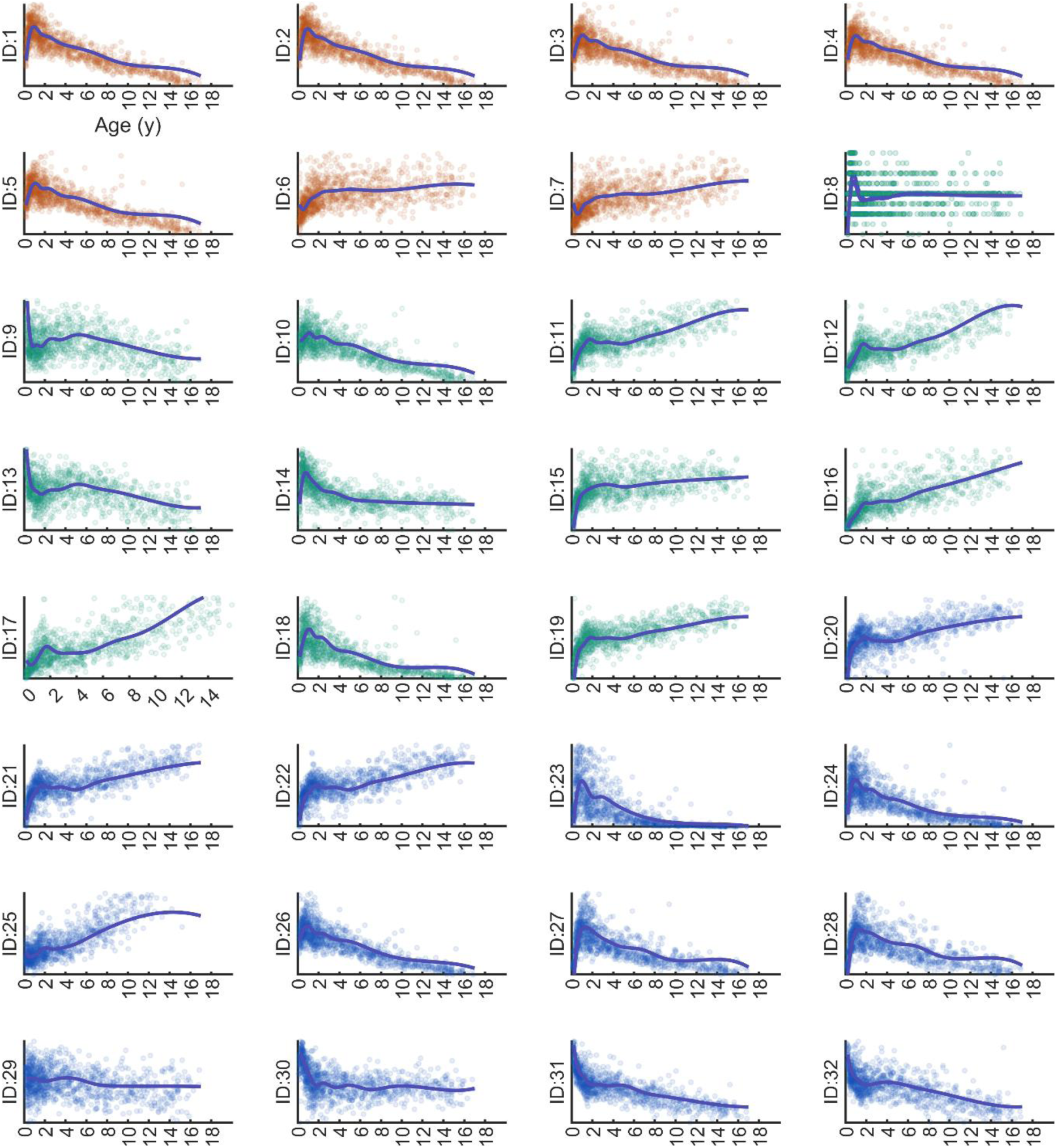
Individual EEG features versus age (all subjects from training dataset D1). As per Supplementary Table 1, each EEG feature and their respective ID are based on 60 epochs from the bipolar montage. Feature types are indicated by colors, where amplitude features (reddish hue), frequency domain features (green hue) and informational metrics (blue hue). A spline is fitted across the median values derived across 2 yearly age bins.

**Supplementary Figure 11.**
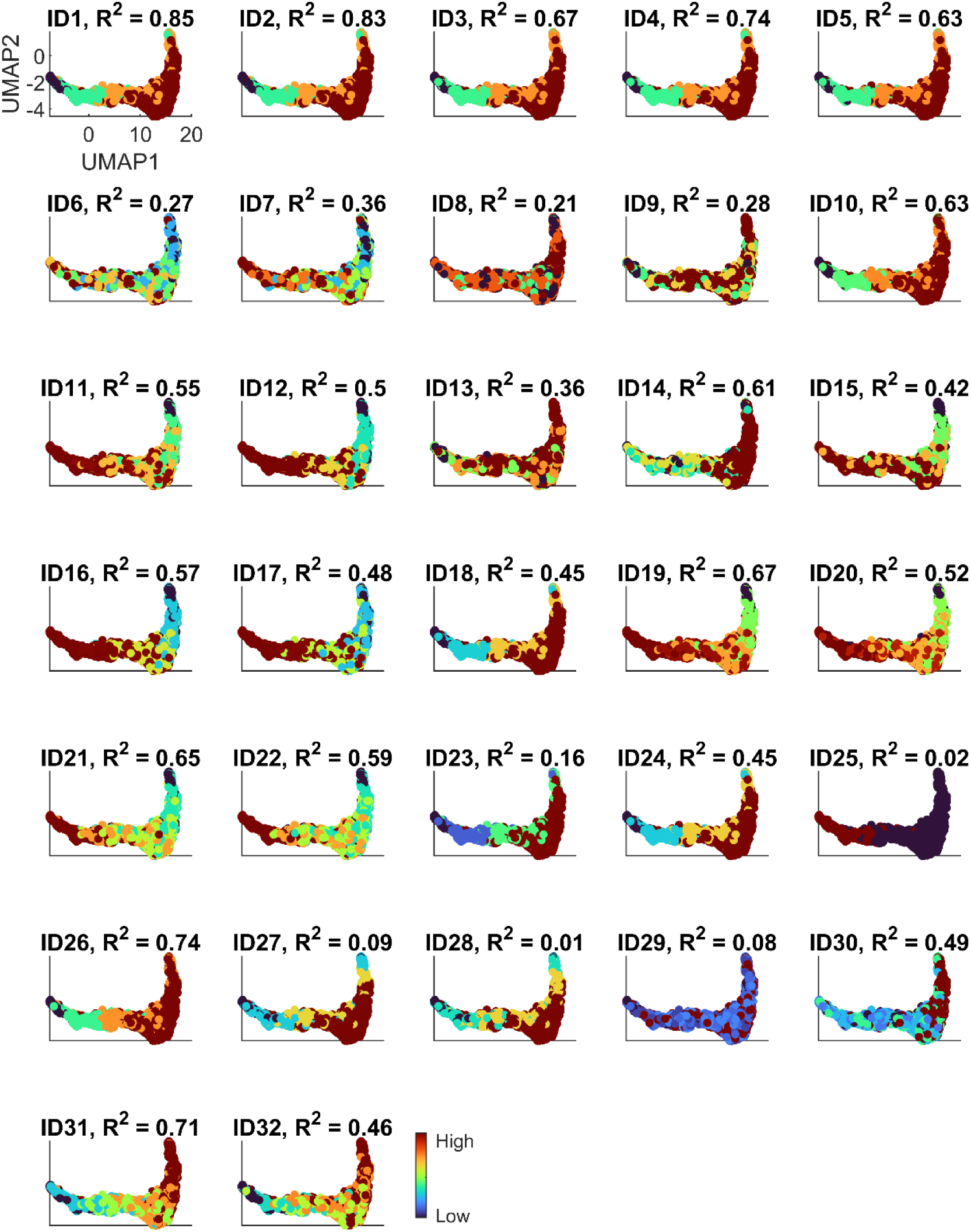
Relating individual EEG features to the late network layer of the Res-NN model. For each individual feature we used GPR to assess the strength of correlation between individual EEG features with network activation layers (here the late layer, i.e., layer 61 from 2-channel EEG data from D1 is shown) to derive a predicted feature value based on UMAP1 and UMAP2 dimensions derived from this training phase. The strength of correlation (R^2^ value) indicates how individual EEG features and predicted individual EEG features are linked by UMAP representations of the Res-NN model, following 10-fold cross-validation. The color bar indicates the relative range of the EEG feature pertaining to its minimum and maximum values (e.g. for feature ID4, low mean amplitude versus high mean amplitude).

**Supplementary Figure 12.**
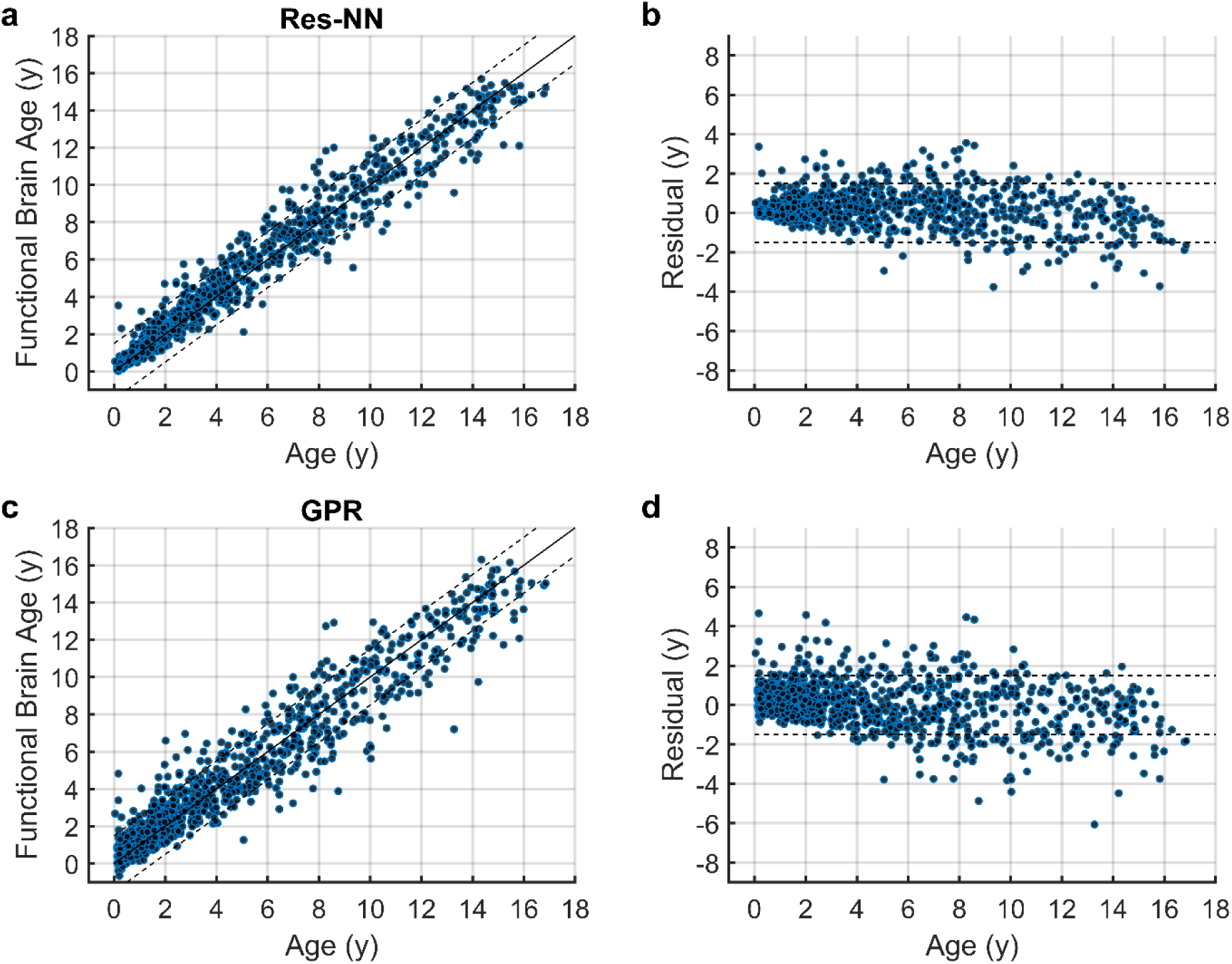
Comparative performance of FBA estimation from a GPR model with the Res-NN. **a.** The Res-NN model (R^2^=0.96, MAE=0.56 years) with error bounds of ±1.5 years and **b.** The residual error (PAD), in years. **c.** GPR results based on a multivariable model of EEG features (R^2^=0.93, MAE=0.79 years). **d.** Residual error (PAD), in years, for the GPR model. Individual EEG recordings (blue filled circles) are plotted across ages.

**Supplementary Figure 13.**
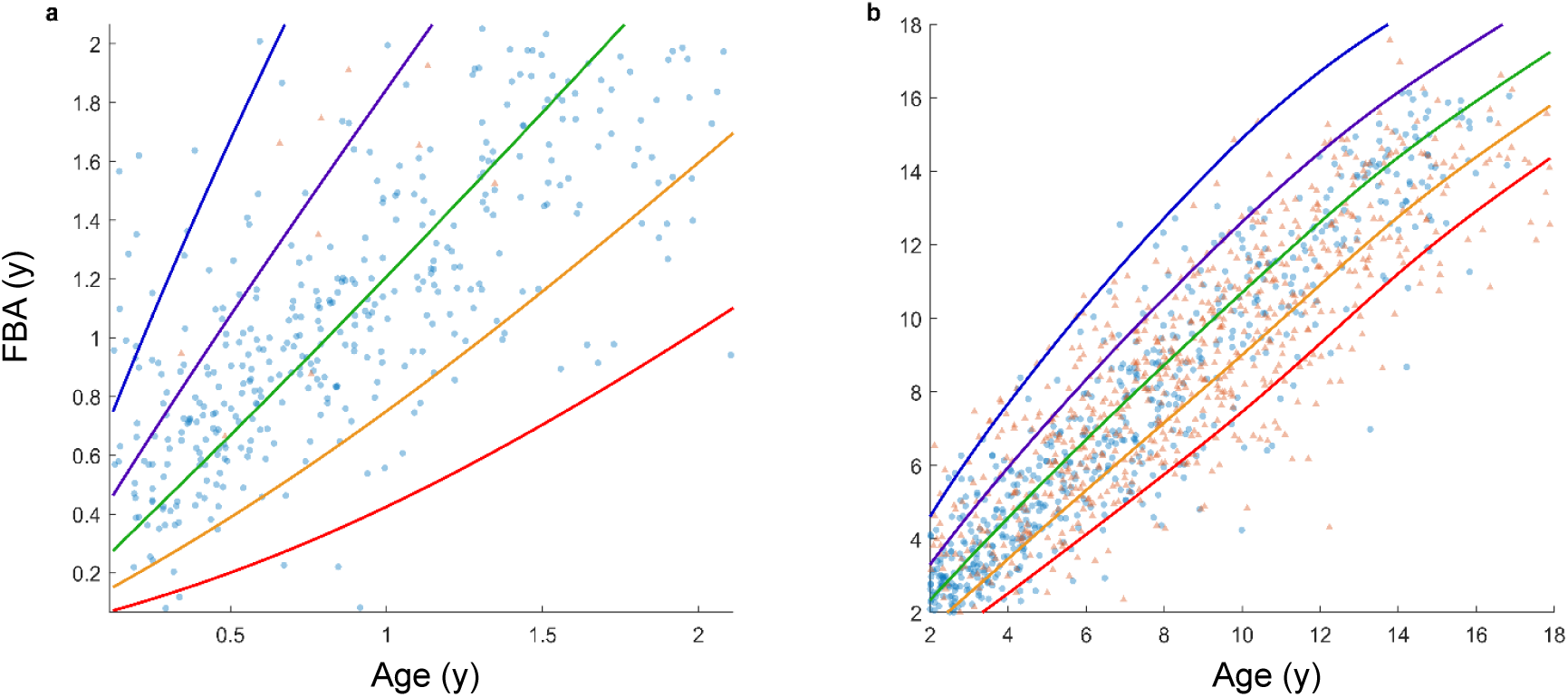
FBA growth charts stratified into infant and childhood age groups. **a.** FBA charts for infants (0 to 2 years) and **b.** children and adolescents (2 to 18 years) were generated to offer a magnified view of the FBA with respect to chronological age. In both age-stratified versions of the chart, over 95% of the data (blue dotted points) are captured between the 3^rd^ and 97^th^ centiles estimated. The 3rd (red), 15th (yellow), 50th (green), 85th (purple) and 97th (blue) centiles are indicated.

**Supplementary Figure 14.**
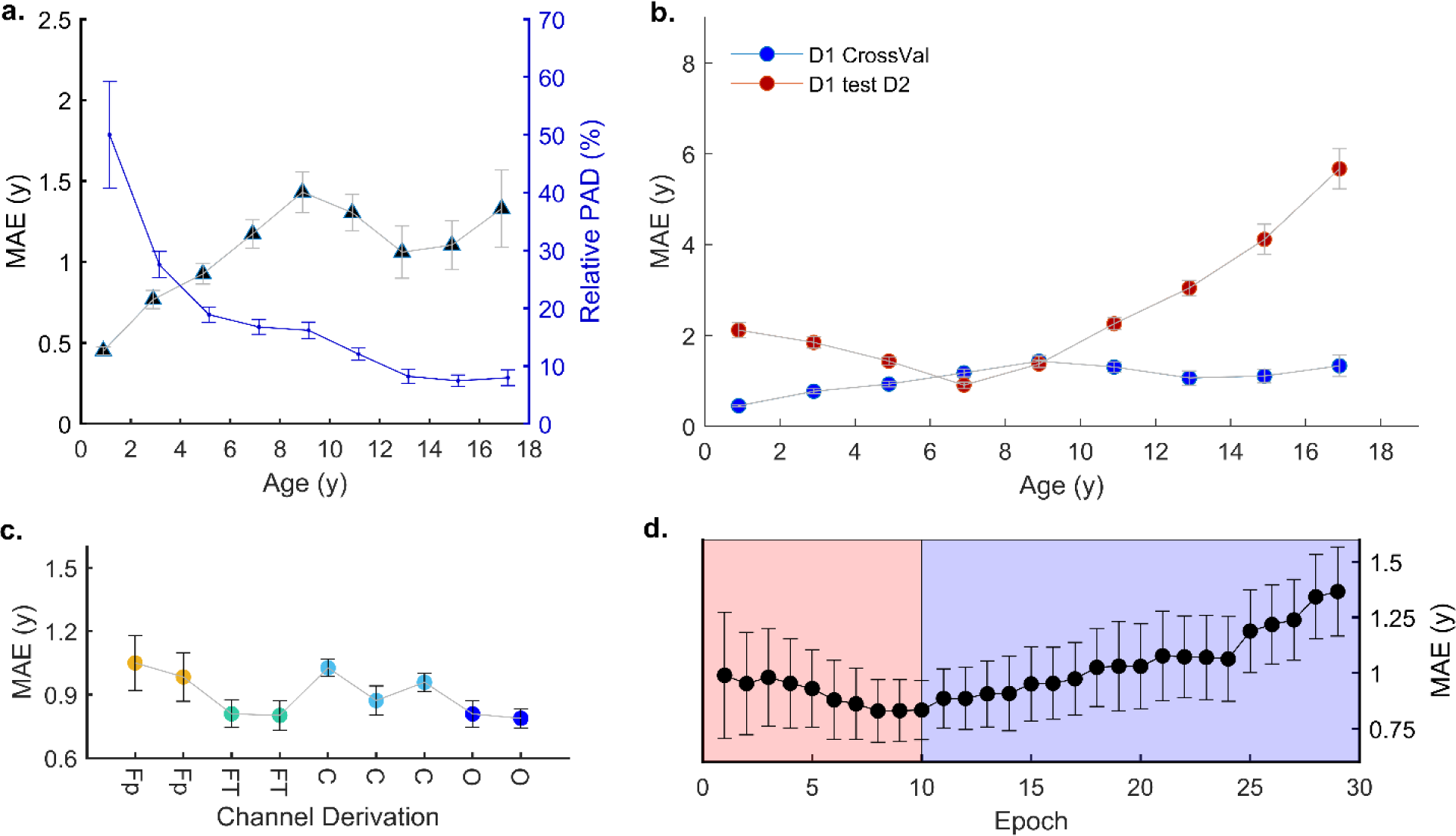
Summary plot of GPR model performance. **a.** The mean absolute error (MAE) of the GPR (dark blue triangles, left axis) across 2 yearly age bins and the relative accuracy of PAD (in blue, right axis) across 2 yearly age bins. For both plots, the mean and standard error of the mean (SEM) are plotted to reflect the sample distribution within age bins. **b.** Differences in MAE of D1 following cross-validation (D1 CrossVal) versus external validation on D2 (D1 test on D2) for GPR. Here differences in sites are more noticeable for younger and older age groups. **c.** Change in MAE across EEG channel locations, with average MAE shown and error bars indicated by standard deviation. **d.** Temporal transitions in the EEG across N1 (light pink) and N2 (light purple). Similar to the Res-NN, the lowest MAE is observed during a transition between sleep N1 and N2 stages. Error bars denote standard deviation for each epoch.

**Supplementary Table 1.** Demographics and additional comparisons of datasets analysed. The key differences between datasets is that EEG in D1 was recorded with a higher channel count as part of outpatient routine clinical EEG whereas EEG in D2 was collected as part of an overnight sleep study PSG. Outcomes in D1 were available at a 4-year follow-up for children in D1 whereas children in D2 had normal sleep study outcome and were hence follow-up outcomes in this group were not available. The age, including the range, median and interquartile range (IQR) are provided along with sex information (and age associated information within sexes).

|  | **D1**<br>Typical Development<br>group (n = 1056) | **D2**<br>Typical Development<br>group (n = 723) | **D2**<br>Trisomy 21 group<br>(n = 40) |
| --- | --- | --- | --- |
| **Age** Range | 6 weeks to 17 years | 3 months to 18 years | 1 year to 17 years |
| Median (years) | 2.7 | 8.1 | 5.4 |
| IQR (years) | 6.2 | 5.6 | 4.2 |
| **Sex**<br>(min, max, median) |  |  |  |
| Males | 543 (6 weeks, 15.8 years, 3 years) | 432 (3 months, 17.9 years, 8.3 years) | 23 (1.2 years, 16.2 years, 4.5 years) |
| Females | 513 (6 weeks, 16.8 years, 2.5 years) | 291 (4 months, 17.8 years, 7.9 years) | 17 (1.9 years, 9.2 years, 5.9 years) |
| **EEG channels available** | 19 | 2 | 2 |
| **Type of clinical EEG** | Outpatient EEG performed in neurology clinic | EEG as part of an overnight sleep study PSG recording | EEG as part of an overnight sleep study PSG recording |
| **Follow-up outcomes** | Yes; neurodevelopmental outcomes were followed up 4 years post-EEG | Not available for this group of children | Not available for this group of children |

**Supplementary Table 2.**
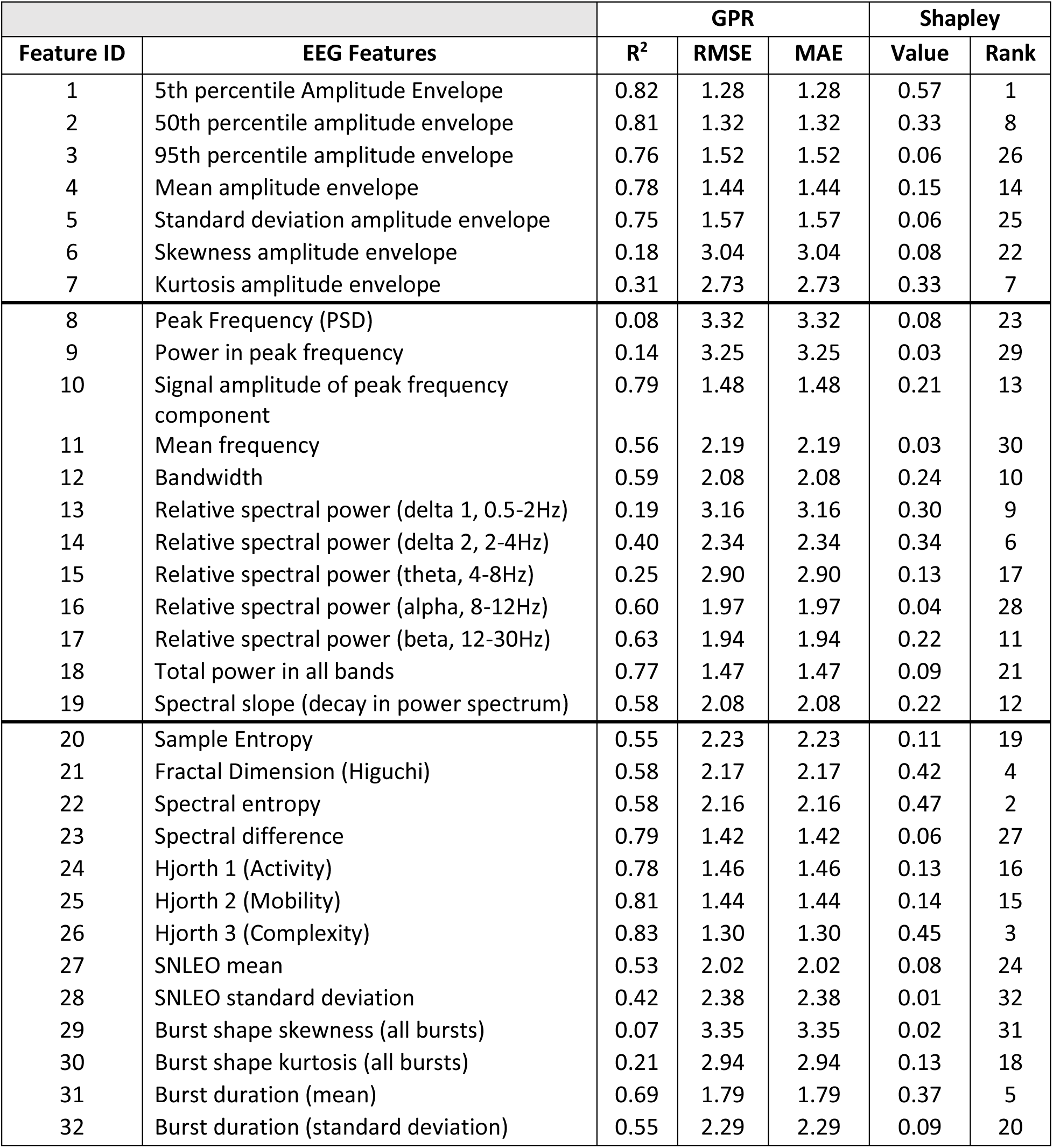
Performance of individual EEG features in predicting age following Gaussian process regression (GPR) and 10-fold cross-validation of training data (D1). Results are based on 60 second EEG epochs acquired in the bipolar montage. Amplitude metrics (ID1 to ID7) denote the 5^th^, 50^th^ (median), 95^th^ centiles based on the absolute value of the Hilbert transform of the EEG (amplitude envelope), along with four central moments of the signal: mean, standard deviation, skewness and kurtosis. Frequency metrics (ID8 to ID19) summarize frequency domain metrics based on the power spectrum density of the EEG signal, and included features such as peak frequency, signal amplitude at this peak, along with mean frequency, bandwidth and relative spectral power in common EEG oscillatory bands. Informational metrics (ID20 to ID32) summarized various non-linear and statistical properties of the EEG. SNLEO stands for the smoothed non-linear energy operator.

**Supplementary Table 3.** Performance of FBA across sleep stages in the validation dataset (D2). The performances of the FBA following 10-fold cross-validation, using either a Res-NN or GPR model, in the D2 dataset. MAE, wMAE and RMSE are shown. N.B. N1 was excluded due to inconsistent availability across ages in D2.

| Sleep stage | # channels, # recordings, # epochs | Res-NN |  | GPR |  |
| --- | --- | --- | --- | --- | --- |
|  |  | MAE (in years) | wMAE (in years) | MAE | wMAE |
| N3 | 2, 723, 13737 | 1.46 | 1.76 | 1.75 | 2.00 |
| REM | 2, 714, 13339 | 1.55 | 1.98 | 1.68 | 1.89 |
| WAKE | 2, 564, 8810 | 1.89 | 2.53 | 2.06 | 2.61 |
| N2 | 2, 723, 13737 | 1.46 | 1.73 | 1.25 | 1.34 |

**Supplementary Table 4.** Overall performance of FBA across datasets. The performances of the FBA, using a GPR model, across all training, test, and combined datasets. MAE, wMAE and RMSE are shown. CI is confidence interval, ^a^only N2 from D1 used, ^b^10-fold cross-validation.

| Train | # channels, # recordings, # epochs | Test | # channels, # recordings, # epochs | R<sup>2</sup> | RMSE (in years; 95% CI) | MAE (in years; 95% CI) | wMAE (in years; 95% CI) |
| --- | --- | --- | --- | --- | --- | --- | --- |
| D1 | 19, 1056, 30624 | D1<sup>b</sup> | 19, 1056, 30624 | 0.93 | 1.09<br>(1.05 – 1.20) | 0.79<br>(0.74 - 0.84) | 1.06<br>(0.85 - 1.27) |
| D1 | 2, 1056, 20064 | D1<sup>b</sup> | 2, 1056, 20064 | 0.91 | 1.24<br>(1.23 - 1.40) | 0.93<br>(0.88 - 0.99) | 1.27<br>(0.97 – 1.57) |
| D1<sup>a</sup> | 2, 1056, 20064 | D2 | 2, 723, 13737 | 0.67 | 2.49<br>(2.34 - 2.64) | 1.94<br>(1.84 – 2.06) | 2.53<br>(2.16 – 2.85) |
| D2 | 2, 723, 13737 | D2<sup>b</sup> | 2, 723, 13737 | 0.84 | 1.41<br>(1.46 - 1.60) | 1.24<br>(1.18 – 1.31) | 1.34<br>(1.12 – 1.57) |
| D1+D2 | 2, 1779, 33801 | D1+D2<sup>b</sup> | 2, 1779, 33801 | 0.90 | 1.43<br>(1.37 – 1.49) | 1.08<br>(1.03 – 1.12) | 1.29<br>(1.12 – 1.47) |

**Supplementary Table 5.** Performance summary across test folds for Res-NN and GPR models. Within each of the 10 test folds, the mean absolute error (MAE) is summarized for both model approaches.

|  | Res-NN | GPR |
| --- | --- | --- |
| Folds | MAE | MAE |
|  | *Test D1* | *Test D1* |
| 1 | 0.61 | 0.72 |
| 2 | 0.49 | 0.73 |
| 3 | 0.59 | 0.73 |
| 4 | 0.53 | 0.78 |
| 5 | 0.59 | 0.75 |
| 6 | 0.49 | 0.76 |
| 7 | 0.50 | 0.98 |
| 8 | 0.63 | 0.80 |
| 9 | 0.59 | 0.86 |
| 10 | 0.55 | 0.80 |
| Fold-wise Average | 0.56 | 0.79 |
|  | *Test D2* | *Test D2* |
| 1 | 1.69 | 1.20 |
| 2 | 1.43 | 1.25 |
| 3 | 1.63 | 1.25 |
| 4 | 1.40 | 1.15 |
| 5 | 1.43 | 1.34 |
| 6 | 1.63 | 1.11 |
| 7 | 1.39 | 1.21 |
| 8 | 1.35 | 1.34 |
| 9 | 1.32 | 1.20 |
| 10 | 1.51 | 1.33 |
| Fold-wise Average | 1.46 | 1.24 |
|  | *Test D1 + D2* | *Test D1 + D2* |
| 1 | 1.15 | 1.07 |
| 2 | 0.97 | 1.05 |
| 3 | 1.03 | 0.91 |
| 4 | 0.95 | 1.14 |
| 5 | 0.96 | 1.21 |
| 6 | 1.06 | 1.16 |
| 7 | 0.89 | 1.11 |
| 8 | 1.36 | 1.04 |
| 9 | 1.00 | 1.13 |
| 10 | 1.11 | 1.03 |
| Fold-wise Average | 1.09 | 1.08 |

**Supplementary Table 6.** Summary of group-wide associations across Res-NN/GPR methods and different cohorts. Assessment of adjusted PAD and centiles derived from the growth chart (Figure 6) where differences between typically developing versus Trisomy 21 (T21) subjects across varying combinations of cohorts (D1, D2 and D1 + D2) are summarized. Effect sizes and differences in adjusted PAD between typically developing children and Trisomy 21 subjects were also examined for subgroups based on sex (i.e. D2 males versus T21 males, D2 females versus T21 females etc). Unpaired t-tests were performed with significance set at p<0.05 with the associated t-statistic reported.

|  | Adjusted PAD |  |  |  | Centiles |
| --- | --- | --- | --- | --- | --- |
|  | Effect size<br>(Cohen's d;<br>95%CI) | Total<br>p-value<br>(t-statistic) | Males<br>p-value<br>(t-statistic) | Females<br>p-value<br>(t-statistic) | p-value<br>(t-statistic) |
| D2 vs T21 | 0.56<br>(0.24 – 0.88) | 0.00053*<br>(3.5) | 0.02*<br>(2.33) | 0.01*<br>(2.56) | 0.028*<br>(3.5) |
| D1 vs T21 | 0.42<br>(0.11 - 0.74) | 0.0087*<br>(2.63) | 0.0012*<br>(3.25) | 0.77<br>(0.29) | 0.08<br>(1.68) |
| D1 + D2 vs T21 | 0.42<br>(0.11 -0.73) | 0.0084*<br>(2.64) | 0.0009*<br>(3.34) | 0.80<br>(0.26) | 0.052<br>(1.92) |
| D2 vs T21 | 0.36<br>(0.06 – 0.68) | 0.023*<br>(2.28) | 0.005*<br>(2.82) | 0.58<br>(0.56) | 0.28<br>(1.08) |
| D1 vs T21 | 0.002<br>(-0.31 - 0.32) | 0.98<br>(0.01) | 0.003*<br>(2.96) | 0.75<br>(0.32) | 0.66<br>(0.44) |
| D1 + D2 vs T21 | 0.16<br>(-0.15 – 0.47) | 0.32<br>(0.98) | 0.31<br>(1.01) | 0.74<br>(0.33) | 0.85<br>(0.19) |

***Supplementary Note 1 -* The effect of data balancing on NN results**

Balancing the dataset is an important component of constructing viable training datasets. In this case, the number of recording per age bin should be equal for the best general regression results; increased diversity of data (typically by adding more data) also provides an improved fit. Furthermore, the training data distribution should best represent the distribution of the larger population. We show the effect of data balancing by training 10 randomly selected network architectures with training datasets containing different levels of heterogeneity with respect to age (***Supplementary Figure 4***). To ensure the number of training points is equal we extract more epochs per recording as the dataset becomes more balanced by changing the overlap of epoch extraction. We select three levels of balance: balanced (uniform distribution of subjects across age) extracts at most 40 recordings from each age group (ages are grouped at yearly intervals), Unbalanced1 (more samples at early ages) extracts at most 80 recordings from each age group, and Unbalanced2 (most samples at early ages) extracts at most 160 recordings from each age group. The RMSE and weighted RMSE (an age adjusted RMSE) are calculated on all EEG recordings that were left out during the 10-fold cross-validation (an unbalanced cohort).

**Supplementary Note 2 – Testing various of network parameters (18 channel FBA for D1)**

We also tested parameters of filter width, filter depth, network depth and filter number within the residual network. The optimal combination was selected using the internal validation data, averaged across all folds. The latter was performed to ensure only 1 network architecture was used across fold. A fixed grid optimization search across was performed across an array of network parameters. For the 18-channel FBA based on dataset D1, this resulted in 180 different network architectures (***Supplementary Figure 6***). The general trends are shown with an optimal combination of network parameters selected: a filter width of 9 samples (*n* = 45 per group), filter depth of 6 channels (*n* = 36 per group), network depth of 3 (*n* = 60 per group), and a filter number of 16 (*n* = 60 per group): a total of 867,601 learnable parameters.

